# Off-target effects of Cre recombinase reveal limits of adoptive T-cell transfers and persistent proliferation of effector CD8 T-cells

**DOI:** 10.1101/502666

**Authors:** Lisa Borkner, Anja Drabig, Xiaoyan Zheng, Julia Drylewicz, Thomas Marandu, Mariona Baliu-Piqué, Suzanne van Duikeren, Ramon Arens, Kiki Tesselaar, José AM Borghans, Luka Cicin-Sain

## Abstract

Effector-memory T-cells (TEM) are assumed to be short-lived cells that poorly proliferate upon antigenic restimulation, thus depending on central-memory T-cells (TCM) to replenish their numbers during homeostasis, largely depending on adoptive transfer evidence. Here we analyzed T cells in their natural environment and observed robust long-term in vivo cycling within the TEM subset that was stronger than the one in the TCM subset. Murine Cytomegalovirus (MCMV) induces inflationary TEM responses that remain high during latency. We analyzed Ki67 expression during acute and latent MCMV infection and found Ki67^hi^Bcl2^lo^ TEM in latently infected mice, arguing for antigen-driven TEM proliferation. TEM acquired deuterium more rapidly than TCM in an in vivo labeling experiment, and were replenished more rapidly than TCM after memory depletion, suggesting that TEM cycle faster than TCM. We depleted selectively the proliferating T-cells by Cre-overinduction, which resulted in a selective loss of Ki67^hi^BCl2^lo^ effector T-cells, and an increase in the death of TEM in the spleen, while it hardly affected the TCM subset, arguing for robust proliferation of TEM in the spleen. On the other hand, TEM homing to the spleen upon adoptive transfer was substantially poorer than TCM, explaining the previously reported expansions of TCM, but not TEM, upon transfer. In conclusion, our data suggest that memory inflation is maintained by proliferation of antigen-specific TEM, rather than by continued expansion and differentiation of TCM.

**Author Summary:** The naïve T cell population consists of T cells that have the potential to recognize millions of different pathogens. Upon infection, naïve T cells that recognize the pathogen expand, and differentiate into effector T cells that eliminate infected cells. Once the infection is contained, the T cell pool contracts and only a small population of central memory T cells remains that can expand quickly upon re-infection. Cytomegaloviruses cause persistent infections that are not cleared from the organism after the initial immune response. In infected individuals a pool of CMV-specific effector memory T cells dominates the immune system in a phenomenon called memory inflation. Previous research using the transfer of central memory or effector memory T cells from CMV-infected mice into mice with a matching infection, showed expansion of central memory T cells but not effector memory T cells. Here we show that effector memory T cells have a reduced capacity to home into lymphoid organs, where T cell activation takes place, compared to central memory T cells. Using methods that do not interfere with T cell differentiation and homing, we show that effector memory T cells are proliferating during the persistent phase of CMV infection, significantly contributing to the upkeep of the inflationary population.

## Introduction

T-lymphocytes play a unique role in the control of intracellular pathogens. T-cell receptor (TCR) recognition of antigenic epitopes presented on MHC molecules results in clonal expansion of activated cells and long-term maintenance of antigen-specific memory cells. This results in natural selection of oligoclonal memory T-cells recognizing previously encountered pathogens, which must be balanced with the need to maintain a broad repertoire of naïve T-cells to allow the recognition of a broad swath of potential infections. Therefore, the proliferation and renewal of the peripheral T-cell compartments is marked by exquisite complexity (1).

The pool of peripheral naïve T-cells is mainly quiescent (2), but ongoing maturation of T-cells from the thymus replenishes it and maintains TCR diversity. The homeostatic turnover of memory CD8 T-cells is more rapid, both in mice and humans (3, 4) and depends on both IL-7 and IL-15 stimulation (5, 6), but occurs independently of T-cell receptor (TCR) stimulation (7). The most rapid proliferation of T-cells is the expansion of CD8 T-cells upon priming in response to antigen (8, 9). This expansion is mainly driven by antigen-specific CD8 T-cells (7), and results in an overall expansion of the effector CD8 T-cells (10). While numbers of effector CD8 T-cells remain elevated by 80 to 160 days post-infection with non-persisting viruses (10), they decline slowly thereafter (11). On the other hand, the expansion of the effector-memory (CD62L^lo^) subset persists for life upon infection with mouse cytomegalovirus (MCMV), a persistent herpesvirus (11).

The human cytomegalovirus (HCMV) latently persists in the vast majority of the adult human population worldwide (12). T-cell responses to HCMV dominate the primed T-cell compartment of seropositive individuals (13). Therefore, CMV-specific T-cells dominate the primed compartment of most adult people in the world, yet the mechanisms of maintenance of these cells remain incompletely understood. Experiments in the mouse model of MCMV infection recapitulated the key aspects of the T-cell response to HCMV infection (14), and allowed the identification of so-called inflationary CD8 T-cell responses – persistence of antigen-specific effector-memory (CD62L^lo^) cells against immunodominant CMV antigens (14–17). It has been proposed that the pool of inflationary T-cells is continuously being replenished by vigorous proliferation of a small subset of central-memory (CD27^+^) T-cells primed early in infection as well as by recruitment of new cells from the naïve T-cell compartment (18). This is in line with the observation that CMV-specific T-cells with effector (CD27^-^) phenotypes show very little proliferative responses upon adoptive transfer into infection matched wild-type mice (19). Furthermore, while inflationary cells are more abundant in lungs than in lymph nodes, the inflationary T-cells expressing the proliferation marker Ki67 were shown to be more abundant in lymph nodes than in lungs (20), suggesting that cycling central-memory T-cells may continuously feed the large pool of inflationary cells in the tissues. On the other hand, blocking lymphocyte egress from lymph nodes does not impair the blood and splenic proliferation of inflationary T-cells (21), and our recent data indicate that inflationary EM cells proliferate by IL-15 driven, homeostatic, proliferation (22). Taken together, these data suggest that the inflationary pool is maintained by systemic hematogenous proliferation of T-cells, rather than by the proliferation of central-memory T-cells in lymph nodes. In conclusion, published data point to scenarios that at face value seem mutually exclusive.

Here we show that Ki67 labeling divides CD8 T-cells into three distinct subsets, where most primed CD8 T-cells are Ki67-intermediate (Ki67^int^) and only the Ki67^hi^ subset represents bona fide cycling cells. The cycling cells in latent MCMV infection displayed predominantly effector phenotypes, whereas cycling was typically homeostatic and restricted to central-memory cells upon clearance of Vaccinia virus. Since adoptively transferred effector-memory and central-memory T-cells were biased in their homing to lymphatic organs, we used alternative methods to validate their contribution to the proliferating pool. In vivo deuterium labeling provided evidence for proliferation of TEM cells in MCMV infection. Selective depletion of primed CD8 T-cells showed that antigen-specific T-cells are rapidly reestablished in MCMV, but not in Vaccinia infection, and that effector-memory T-cells rebound before central-memory T-cells. Targeted depletion of cycling cells significantly increased the apoptosis of effector-memory T-cells in the spleen and decreased the count of inflationary T-cells, while leaving the central-memory T-cells hardly affected, substantiating the claim that inflationary CD8 T-cells maintain their numbers by antigen-driven cycling of effector-memory T-cells. Taken together our data indicate that terminally differentiated T-cells retain a substantial cycling potential and that the continuous cycling of effector T-cells is critical for the maintenance of inflationary responses.

## Results

### Ki67 exhibits three distinct expression levels in CD8 T-cell populations during acute and long-term responses to infection

To define the amount of T-cells proliferating in the memory phase of a persistent or a resolved infection, we infected mice with MCMV or the non-persistent Vaccinia virus (VACV) and analyzed blood CD8 T-cells for the expression of cycling marker Ki67 and the DNA marker FXcycle at 120 days postinfection (dpi). We used as controls the blood from the very same mice during the acute phase, at 7 dpi.

To our surprise, the intracellular staining of Ki67 did not simply result in a Ki67 positive and a negative population, but in three distinct populations, which we will further on call Ki67^-^, Ki67^int^, and Ki67^hi^. Cells within the S, G2, or M phase as indicated by an increased amount of DNA (FXcycle^+^) were predominantly found in the Ki67^hi^ population at 7 dpi (Fig. 1A, Supplementary Fig. 1A), but some were also encountered in the Ki67^int^ fraction, and very few in the Ki67^-^ subset (Fig. 1A).

**Figure 1:**
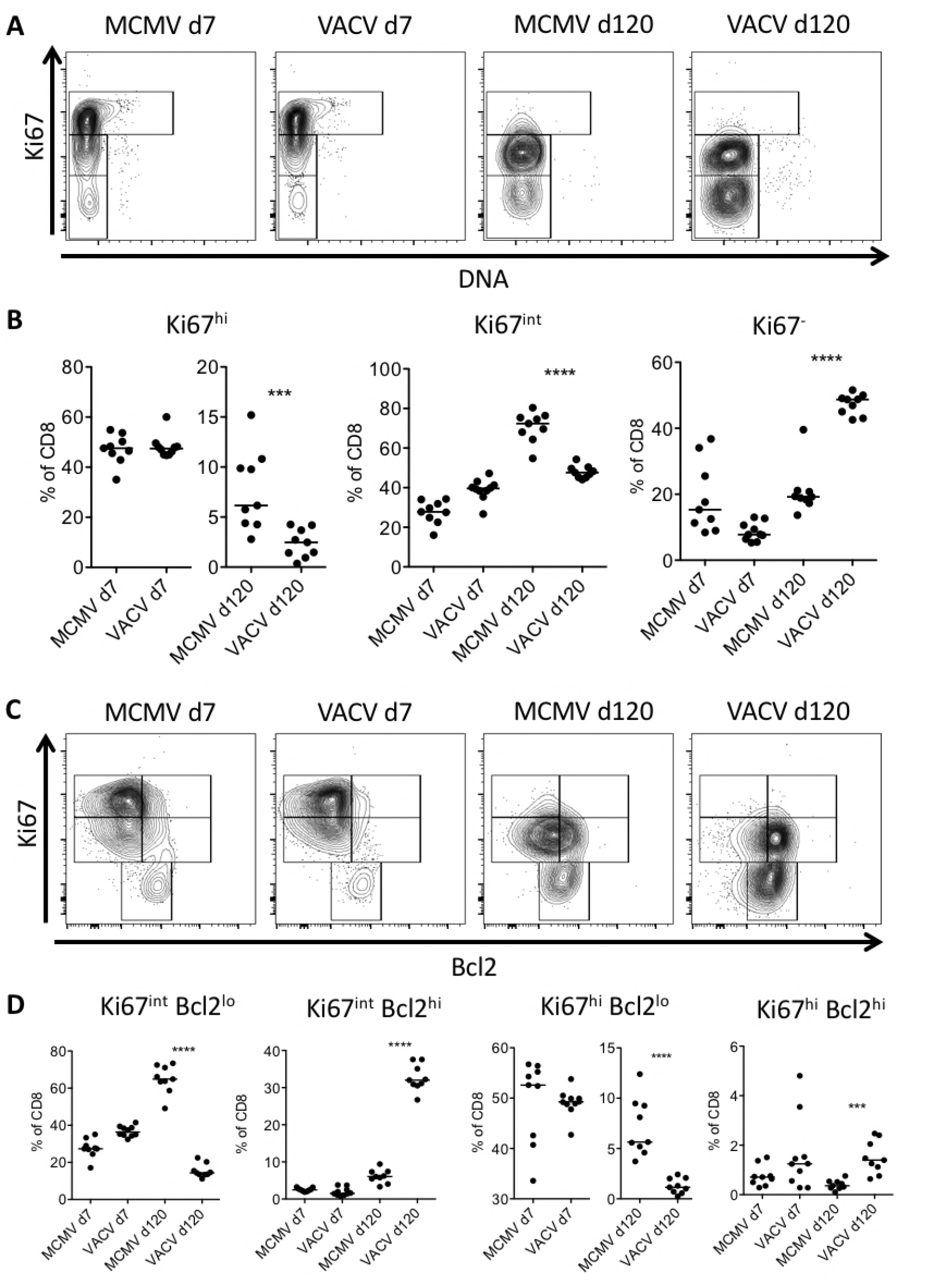
Latently MCMV infected mice preserve antigen-driven proliferation of T-cells. 129/Sv mice were infected with 2 × 10^5^ PFU MCMV or VACV. At 7 and 120 dpi, blood was taken and proliferating T-cell subsets were analyzed with antibodies directed against CD8, Ki67 and Bcl2. In addition, the DNA content of the cells was determined using the DNA marker FXcycle violet. (A) Representative dot plots demonstrating the gating of cell cycle subsets based on Ki67 and FXcycle violet. (B) Percentage of cell cycle subsets in total CD8 T-cells. (C) Representative dot plots demonstrating the gating of cell cycle subsets based on Ki67 and Bcl2. (D) Percentage of Ki67 Bcl2 subsets in total CD8. B and D: each dot represents one mouse; horizontal lines indicate median. Mann-Whitney Test was used to assess significance at **=p<0.01, ***=p<0.001 and ****=p<0.0001.

At 7 dpi, the fraction of Ki67^-^, Ki67^int^, and Ki67^hi^ subsets of CD8 T-cells were roughly similar in MCMV and VACV infected animals. Approximately 50% of cells were observed in the Ki67^hi^ population, 30-40% in the Ki67^int^, and less than 20% of CD8 T-cells in the Ki67^-^ fraction, consistent with the strong antigen-driven proliferation of CD8 T-cells that one would expect during the acute phase of viral infection (Fig. 1B). At 120 dpi, this picture changed considerably: In both infections, the percentage of Ki67^hi^ CD8 T-cells was substantially reduced, but it remained significantly higher in MCMV infected mice (median of 6.2%) than in VACV infected mice (median of 2.5%). The Ki67^int^ population remained at around 40% in the VACV infection, but made up as much as 70% of the CD8 T-cell compartment in MCMV infected mice. The fraction of Ki67^-^ cells showed a median value close to 20% in the MCMV-infected group but approximately 50% in VACV infected mice and was thus significantly larger in VACV infected mice (Fig. 1B). Assuming that at least Ki67^hi^ cells are cycling, our data indicated that the robust cycling of CD8 T-cells upon acute infection is followed by quiescence in VACV infected mice and by continuous cycling at a low level in latent MCMV infection at 120 dpi. It remains unclear if Ki67^int^ cells were also a cycling subset, which would have implied an even larger difference in CD8 T-cell cycling at 120 days post MCMV or VACV infection.

A key difference in T-cell cycling during acute or resolved infections is that cycling is antigen-driven in acute infection and homeostatic after resolution. Therefore, to better understand the difference between Ki67^hi^ and Ki67^int^ expression, we analyzed the Ki67 subsets of CD8 T-cells for the expression of Bcl2, an anti-apoptotic gene that is commonly used as marker of homeostatic proliferation and that is down-regulated after TCR engagement and T-cell activation (23, 24). We found that the markers Ki67 and Bcl2 determine five different populations in the CD8 T-cell compartment: non-cycling Ki67^-^ T-cells, Ki67^int^ Bcl2^lo^, Ki67^int^ Bcl2^hi^, Ki67^hi^ Bcl2^lo^, and Ki67^hi^ Bcl2^hi^ T-cells (Fig. 1C).

The pattern of Ki67 and Bcl2 expression in MCMV and VACV infection was comparable at 7 dpi, with most cells in the Ki67^int^Bcl2^lo^ or Ki67^hi^Bcl2^lo^ populations, indicating antigen-driven proliferation (Fig. 1C, D), but starkly different at 120 dpi. Here, the most abundant fraction in MCMV infected mice was Ki67^int^Bcl2^lo^, whereas the Ki67^int^ cells in VACV infected mice were mainly Bcl2^hi^ and this difference was significant (Fig. 1C, D). Similarly, significantly higher percentages of Ki67^hi^Bcl2^lo^ CD8 T-cells were observed in MCMV infected mice (Fig. 1C, D) than in the VACV group (median 5.6% versus 1.1%, respectively). Taken together, our results argue that CD8 T-cells expressing Ki67 in resolved VACV infection fit the pattern of homeostatic proliferation, but most cells in latent/persistent MCMV infection do not.

### Cycling CD8 T-cells in latent MCMV infection are largely effector T cells

We considered it likely that differences in Bcl2 expression reflect antigen-driven proliferation in latent MCMV infection versus homeostatic proliferation in the resolved VACV infection. We reported previously that primed T-cells assume a memory phenotype upon VACV infection over time, whereas latent MCMV infection sustains a large pool of effector cells (11), which are known to lack substantial Bcl2 expression. To determine if the difference in Ki67 and Bcl2 expression upon MCMV and VACV infection reflects a difference in the phenotype of cycling cells, Ki67^hi^ and Ki67^int^ cells were analyzed for the expression of the surface markers CD44 and CD127, allowing us to separate CD44^hi^CD127^hi^ memory T-cells (TM), CD44^hi^CD127^lo^ effector T-cells (TE), and CD44^lo^CD127^hi^ naïve T-cells (Fig. 2A). Consistent with previous reports (11), effector T-cells were more frequent in MCMV than in VACV infection at 120 dpi and displayed a Bcl2^lo^ phenotype (data not shown). Ki67^-^ cells showed a predominantly naïve phenotype in both groups (data not shown), and were not analyzed further. On the other hand, both the Ki67^hi^ and the Ki67^int^ subset displayed a significantly higher frequency of TE cells in MCMV-infected mice, and conversely a higher frequency of TM cells in the VACV infected group (Fig. 2B). Therefore, we concluded that the significantly higher frequency of Ki67^hi^ and Ki67^int^ CD8 T-cells in MCMV-infected mice at day 120 was mainly due to the high frequencies of Bcl2^lo^ effector T-cells found in long-term MCMV infection.

**Figure 2:**
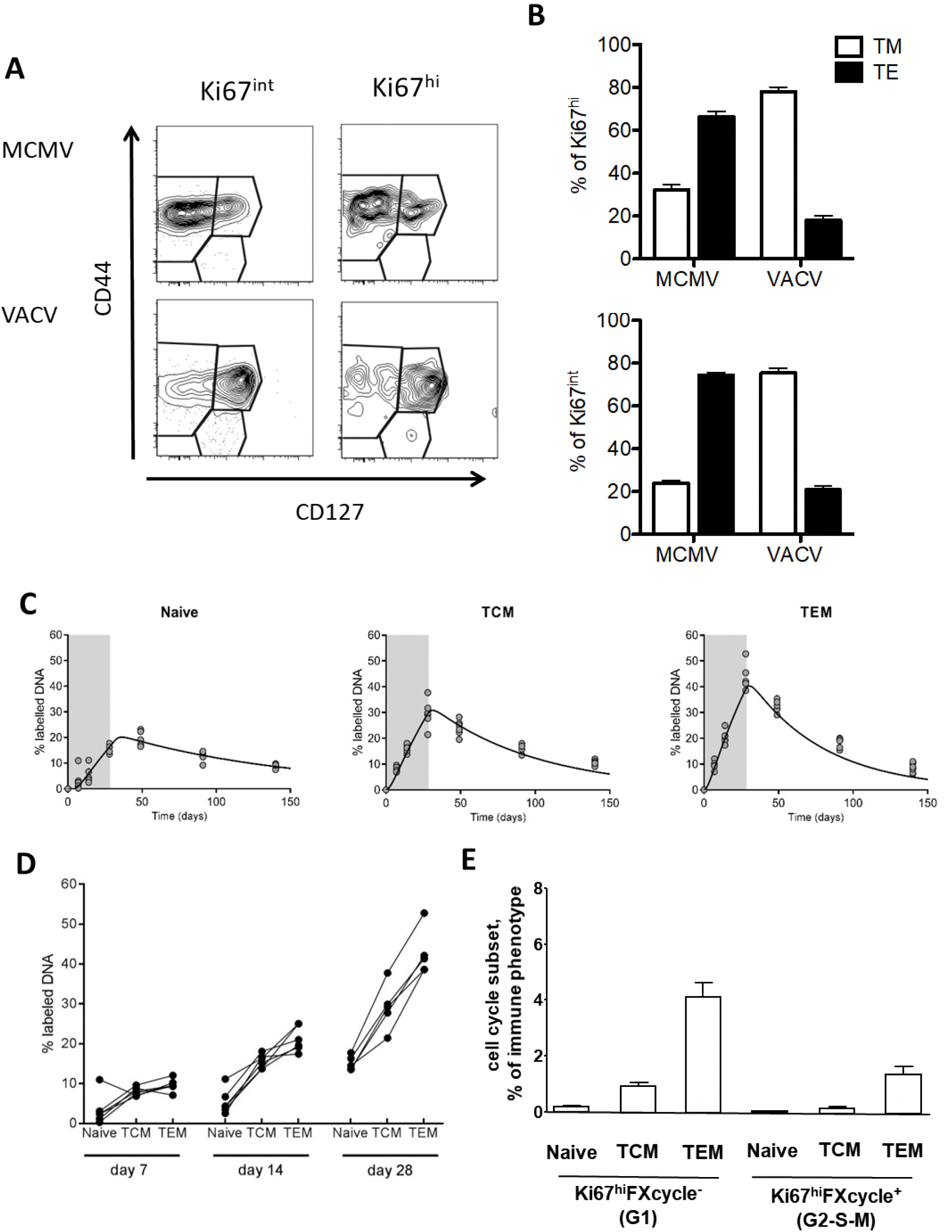
Proliferating CD8 T-cells in latently MCMV infected mice are largely effector cells. (A) and (B): 129/Sv mice were infected with 2 × 10^5^ PFU MCMV or VACV. At 7 and 120 dpi, blood was taken and proliferating T-cell subsets were analyzed with antibodies directed against CD8, Ki67, CD44, and CD127. (A) Representative dot plots demonstrating the determination of the immune phenotype using CD44 and CD127. (B) Percentage of TM (CD44^+^ CD127^+^) and TE (CD44^+^ CD127^-^) in cycling (Ki67^int^ and Ki67^hi^) subsets of CD8 T-cells. Bars indicate means, error bars indicate SEM. (C) and (D): 129/Sv mice were infected with 2 × 10^5^ PFU MCMV. From 120 dpi onward they were given D_2_O for 28 days. Mice were sacrificed at indicated time points and CD8^+^ tetramer-negative splenocytes were sorted on CD44 and CD62L expression to isolate DNA from TEM (CD62L^-^ CD44^+^), TCM (CD62L^+^ CD44^+^), and naïve (CD62L^+^ CD44^-^) CD8 T-cells. The deuterium content in the DNA of these CD8 T-cell subsets was determined by mass spectrometry. Three independent experiments with 5-6 mice per time-point were performed, and data from a representative experiment are shown. (C) Best fits of the mathematical model (solid curves) to the deuterium enrichment levels in the indicated cell subsets(circles for individual mice). The gray-shaded area indicates the window of D_2_O administration. Parameters corresponding to the best fits are summarized in Table 1. (D) Comparison of the percentage of deuterium labelled DNA in the three T-cell subsets of each mouse at the indicated time points during label administration. Each symbol represents a T-cell subset from one mouse and lines connect data points from the same mouse. (E) Blood leukocytes from C57BL/6 mice gated by CD62L and CD44 expression into naïve, TCM, and TEM subsets at 100 dpi with MCMV were labelled intracellularly with Ki67 and FXcycle (see Fig 3A for representative gating) and mean frequencies of Ki67^hi^ populations in indicated subsets are shown. Experiments were performed twice (total n=12). Error bars are SEM.

### Effector CD8 T-cells proliferate in latently MCMV infected mice

We independently quantified CD8 T-cell proliferation in latently MCMV-infected mice by in vivo stable isotope labeling, a method that is not toxic and does not interfere with cell dynamics. Mice were given deuterated water (D_2_O) at 120 dpi over a course of 4 weeks.

We measured deuterium levels in the DNA of different CD8 T-cell subsets during the labeling period and for additional 16 weeks thereupon. We sorted CD8^+^ splenocytes into CD44^+^CD62L^+^ (central-memory – TCM), CD44^+^CD62L^-^ (effector-memory – TEM) and naïve (CD44^-^CD62L^+^) T-cells at various days during and after label administration, isolated DNA from the sorted subsets, and quantified the fraction of deuterium labeled DNA. Pairwise comparison of the level of deuterium enrichment in the different T-cell subsets revealed that at every time point during up-labeling, deuterium enrichment was lowest in naïve cells and consistently higher in TEM than in TCM cells (Fig. 2D), arguing for robust TEM proliferation. Using mathematical modeling (See material and methods), we estimated the average turnover rate *(p)* of TCM, TEM and naïve T-cells, i.e. the fraction of cells replaced by new cells per day, and their corresponding lifespans *(1/p)*. The best fits to the data (Fig. 2c) suggested that naïve T-cells had a relatively low turnover rate of 0.8% per day, TCM cells an almost two-fold higher turnover rate of 1.4% per day, but TEM cells displayed the highest turnover rate of 2.0% per day (see Table 1). Although part of the label incorporation in the TEM population may have been obtained during T-cell proliferation at the TCM stage after which these cells differentiated into TEM cells (25), the significantly higher turnover rate and consistently higher level of deuterium enrichment in TEM cells shows that TEM cells themselves also proliferate significantly.

**Table 1:** Parameters corresponding to the best fits of the mathematical model to the deuterium labeling data.

|  | Naïve | TCM | TEM |
| --- | --- | --- | --- |
| turnover rate $p$ (per day) | 0.0083<br>[0.0070; 0.0088] | 0.0138<br>[0.0119 ; 0.0141] | 0.0196<br>[0.0169 ; 0.0199] |
| Average lifespan ($1/p$ in days) | 120 [113;142] | 72 [71;84] | 51 [50;59] |

\*values in brackets represent 95% confidence intervals on the parameters.

Finally, to validate independently the rate of cycling in the naïve, TCM, and TEM subsets in another mouse strain, blood leukocytes from latently infected C57BL/6 mice were assessed by Ki67 and FXcycle staining. Ki67^hi^ FXcycle^-^ cells were considered as cycling cells in the G1 phase, whereas Ki67^hi^FXcycle^+^ cells were assumed to be in the G2, the S, or the M phase of the cell cycle. The percentages of Ki67^hi^ G1 and Ki67^hi^ G2-S-M cells were clearly higher in the TEM compartment than in the TCM compartment, while the naïve subset showed very few cycling cells (Fig. 2E). Taken together, these data argue for mouse-strain independent TEM proliferation in latently MCMV-infected mice.

### Activation of Cre-recombinase in R26 CreER^T2^ mice leads to specific loss of proliferating CD8 T-cell subsets in blood and spleen

While D_2_O labeling demonstrated proliferation and turnover of primed T-cell subsets in MCMV latency, and Ki67^hi^ cells were most abundant among effector-memory T-cells, it remained unresolved if Ki67^hi^, Ki67^int^, or both subsets are the bona fide cycling ones and replenish the CD8 compartment. To specifically test the proliferation of Ki67^hi^ and Ki67^int^ CD8 T-cells we designed an assay that targets cycling cells for depletion. Persistent overexpression of Cre-recombinase leads to non-targeted effects beyond loxP sites and illegitimate chromosomal recombination (26). This effect disproportionately affects the rapidly proliferating cells in the hematopoietic system, resulting in catastrophic chromosomal aberrations in the dividing cells and thus in their death and loss (27). To target dividing cells for depletion once MCMV had established latency, we used the Rosa26-CreER^T2^ mice (R26 CreER^T2^), which ubiquitously express a variant of the Cre-recombinase that can be activated by the administration of Tamoxifen (Tam). Latently infected R26 CreER^T2^ mice were administered Tam for five consecutive days followed by a three day break and one more day of Tam, upon which we analyzed the proliferation of blood lymphocytes and splenocytes by flow cytometry. Using the markers Ki67 and FXcycle we distinguished four subsets: Ki67^-^, Ki67^int^, Ki67^hi^FXcycle^-^, and Ki67^hi^FXcycle^+^ (Fig. 3A). In absence of Tam, approximately 2% of CD8 T-cells were Ki67^hi^FXcycle^-^ in both blood and spleen, and even fewer cells were seen in the Ki67^hi^FXcycle^+^ subset (Fig. 3A and 3B). Hence, both compartments showed some ongoing CD8 T-cell proliferation in latent MCMV infection. Mice that were treated with Tam displayed a clear loss of FXcycle^+^ cells over controls receiving vehicle only (Fig. 3A), demonstrating targeted depletion of CD8 T-cells in the S, G_2_, or M phase of the cell cycle by Tam treatment. Ki67^hi^FXcycle^-^ cells were also depleted (Fig. 3A), arguing that this subset is targeted by the same mechanism. This resulted in significantly reduced Ki67^hi^Fxcycle^-^ and Ki67^hi^Fxcycle^+^ populations (Fig. 3B). On the other hand, we observed no significant loss of Ki67^int^ cells, and a mild increase in the percentage of Ki67^-^ cells (Fig. 3B). Since Tamoxifen treatment reduced specifically the CD8 T-cell subsets with Ki67^hi^ expression, but had no effect on Ki67^int^ cells, our data suggested active cycling of Ki67^hi^ CD8 T-cells and quiescence in most of the Ki67^int^ population (Fig. 3B).

**Figure 3:**
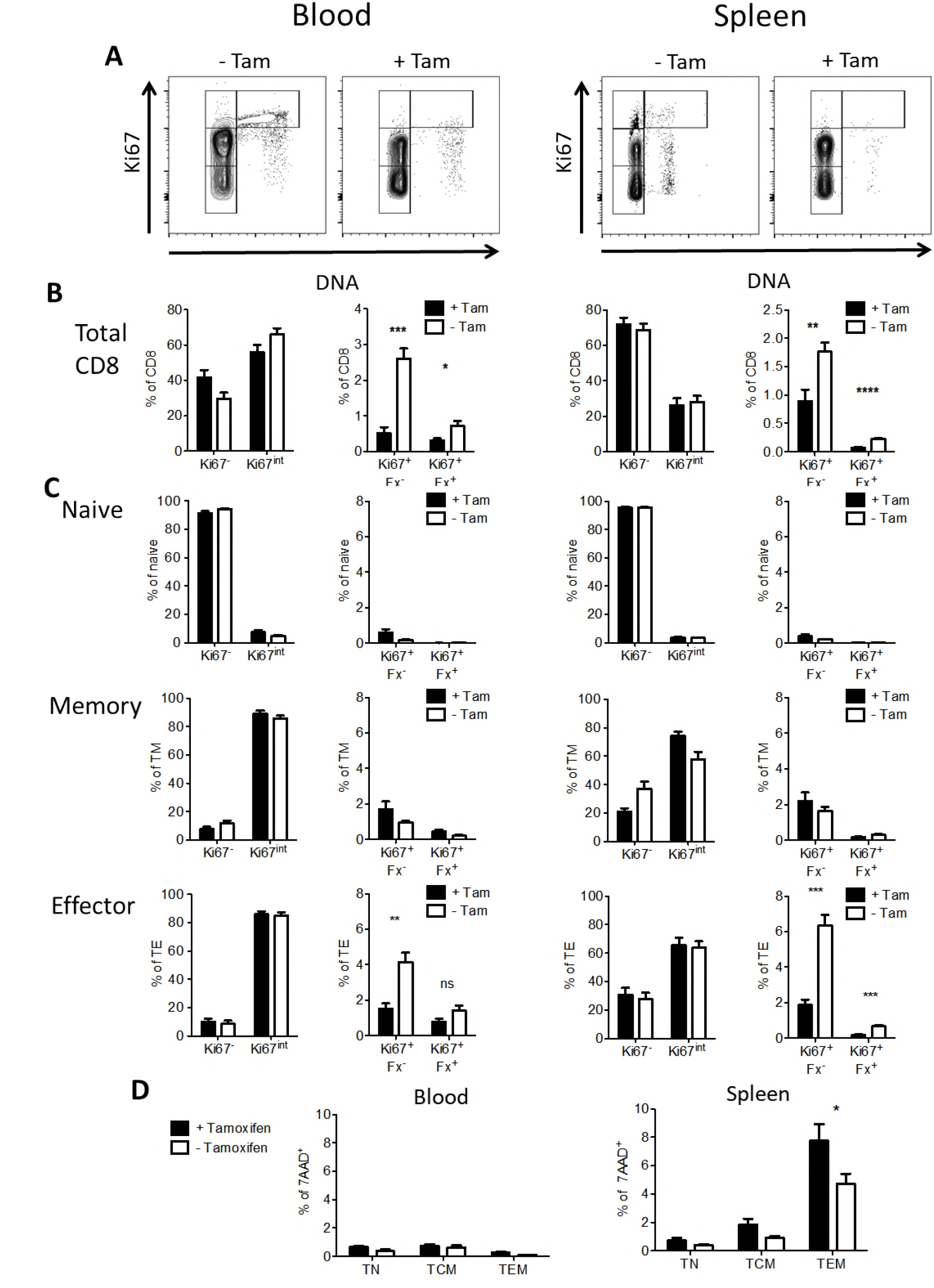
The activated Cre-Recombinase induces cell death in Ki67^hi^ effector T-cells in blood and spleen. R26CreER^T2^ mice were infected with 10^6^ PFU MCMV. At 94 dpi the mice received 80mg/kg BW Tamoxifen by oral gavage for 5 consecutive days, followed by a three day break, and one more day of treatment, before mice were sacrificed. Cell proliferation in CD8 T-cell subsets was characterized by CD8, CD44, CD127, and Ki67 expression as well as by the DNA marker FXcycle. (A) Representative dot plots for CD8 T-cell proliferation in blood and splenocytes in treated and untreated animals. (B) Percentages of cell cycle subsets Ki67^-^, Ki67^int^, Ki67^hi^FX^-^, and Ki67^hi^ FX^+^ in total CD8 T-cells. (C) Percentages of cell cycle subsets Ki67^-^, Ki67^int^, Ki67^hi^FX^-^, and Ki67^hi^ FX^+^ in naïve (CD127^+^ CD44^-^), memory (CD127^+^ CD44^+^), and effector (CD127^-^ CD44^+^) CD8 T-cell subsets. (D) The percentage of dead cells (7-AAD^+^) was determined in the naïve (CD62L^+^ CD44^-^), central-memory (CD62L^+^ CD44^+^), and effector-memory (CD62L^-^ CD44^+^) CD8 T-cell subsets in blood and spleen. Pooled data from two independent experiments (12 mice in total) are shown. Bars indicate means, error bars indicate SEM. Significant differences were determined by Mann-Whitney test, *=p<0.05, **=p<0.01, ***=p<0.001, and ****=p<0.0001

### Tamoxifen targets splenic effector T-cells for depletion

Both the Ki67 staining and the deuterium labeling results were consistent with robust proliferation of effector (CD127^-^) or effector-memory (CD62L^-^ CD127^+/-^) T-cells. Tam treatment affected the relative size of blood and splenic T-cell subsets, decreasing the size of the effector subset (CD127^-^) and increasing relatively all the other ones (Suppl. Fig. 2A and data not shown). However, these data did not exclude a scenario in which rapidly cycling central-memory or effector-memory T-cells in MCMV infection downregulate CD62L and CD127 and assume an effector phenotype. A major contribution of central-memory T-cells to the maintenance of inflationary responses was previously suggested by adoptive transfer experiments (19). In that case, memory cells, rather than effector cells, would be the major driver of cycling in MCMV latency. Hence, we analyzed the Ki67 and FXcycle staining patterns in the effector (CD127^-^ CD44+), memory (CD127^+^ CD44^+^), and naïve (CD127^+^ CD44^-^) CD8 T-cells of R26 CreER^T2^ mice after Tam treatment.

In naïve T-cells, both in blood and spleen, over 90% of cells were in the Ki67^-^ gate and Tam treatment had no discernible effect on this subset. Most memory CD8 T-cells in blood and spleen were in the Ki67^int^ population. This subset also showed no significant change in the cell fractions upon Tam depletion (Fig. 3C). Effector CD8 T-cells were also observed predominantly in the Ki67^int^ population, but this subset also showed the highest fraction of cells in the Ki67^hi^ gates (both FXcycle^+^ and FXcycle^-^) in absence of Tam. A highly significant reduction of Ki67^hi^FXcycle^+^ and Ki67^hi^FXcycle^-^ fractions could be observed in the spleen upon Tam treatment. The percentage of Ki67^hi^ T-cells was also reduced in the Ki67^hi^FXcycle^-^ subset in the blood (Fig. 3C), implying that targeted depletion of cycling cells affected disproportionately the effector subset in both compartments. Similarly, gating on CD62L and CD127 expression revealed low levels of Ki67 and FXcycle labeling in CD62L^-^CD127^+^ T cells and a pronounced reduction of Ki67^hi^ subsets upon Tam-treatment only in the CD62L^-^CD127^-^ subset (data not shown).

The reduction of Ki67^hi^ subsets in blood and spleen indicated that effector CD8 T-cells may cycle in either compartment, but did not allow us to define if cycling is restricted to one compartment over the other. We reasoned that a more immediate assay to define the cycling activity would be to measure cell death upon Tam treatment. Therefore, we stained the CD8 T-cells of latently infected R26 CreER^T2^ mice with antibodies against CD62L and CD44 and determined the fraction of dead cells via 7AAD. We observed very few 7AAD^+^ lymphocytes in the blood, and a substantially higher percentage of 7AAD^+^ cells after Tam treatment in the spleen (Fig. 3D). This increase was particularly pronounced in the TEM subset of CD8 T-cells (Fig. 3D) and similar results were observed using Annexin V staining (data not shown). R26 CreER^T2^ mice infected with MCMV for a very long time (21 months) prior to Tam treatment and administered with Tam-spiked food pellets for two weeks showed a similar significant increase of 7AAD^+^ cells only in the TEM fraction of splenic CD8 T-cells (Supplementary Fig. 2B). Hence, cycling of effector-memory T-cells in spleens of latently infected hosts is maintained essentially for life. We also analyzed the expression pattern of Ki67 and Bcl2 in CD8 T-cells of R26 CreER^T2^ mice treated with Tam. Using the gating scheme introduced in Fig. 1C, we found a robust population of Ki67^hi^Bcl2^lo^ CD8 T-cells in the blood and in the spleen of untreated mice (Supplementary Fig. 2C), indicating non-homeostatic proliferation, likely due to ongoing antigen stimulation. This population, however, disappeared almost completely after Tam treatment and this decrease was specific for this gate and highly significant in both blood and spleen (Supplementary Fig. 2D).

Taken together, our data show that Ki67^hi^ effector cells are more abundant and more susceptible to Tam treatment in R26 CreER^T2^ mice than Ki67^hi^ naïve or memory T-cells, particularly in the spleen, thereby identifying Ki67^hi^ effector cells as main proliferating cells. The observed Cre-recombinase effects are most pronounced in Bcl2^-^ cells arguing for a robust antigen-driven T-cell proliferation.

### Adoptive transfer of TCM and TEM leads to skewed distribution of T-cells to secondary lymphoid organs

Our deuterium labeling and depletion data argued for robust cycling of effector T-cells, which was unexpected because studies using adoptive transfer suggested that inflationary effector CD8 T-cells can only be replenished from a pool of central-memory T-cells (19, 20). We speculated that the discrepancy might be due to differences in the experimental design. Since we recently observed an association between the latent MCMV load in the spleen and the size of memory inflation (28) and since Tam induced apoptosis in splenic TEM cells (Fig. 3D), we considered it likely that the spleen is a major cycling site. Consequently, adoptive transfer would skew the experimental results towards T-cell subsets that efficiently home to lymphatic organs upon transfer. To test this idea, we sorted CD62L^+^CD44^+^ or CD62L^-^ CD44^+^ CD8 T-cells from splenocytes of latently infected Ly5.1 C57BL/6J mice and transferred them into naïve Ly5.2 C57BL/6J mice. 18 hours after adoptive transfer, we found significantly more CD62L^+^CD44^+^ than CD62L^-^CD44^+^ CD8 T-cells in the spleen (Fig. 4A). Similarly, antigen-specific CD62L^+^ cells homed to the spleen more efficiently than their CD62L^-^ counterparts (Supplementary Fig. 3). In consequence, adoptive transfer skews experimental results towards the CD62L^+^ cell subset, which efficiently homes to the spleen.

**Figure 4:**
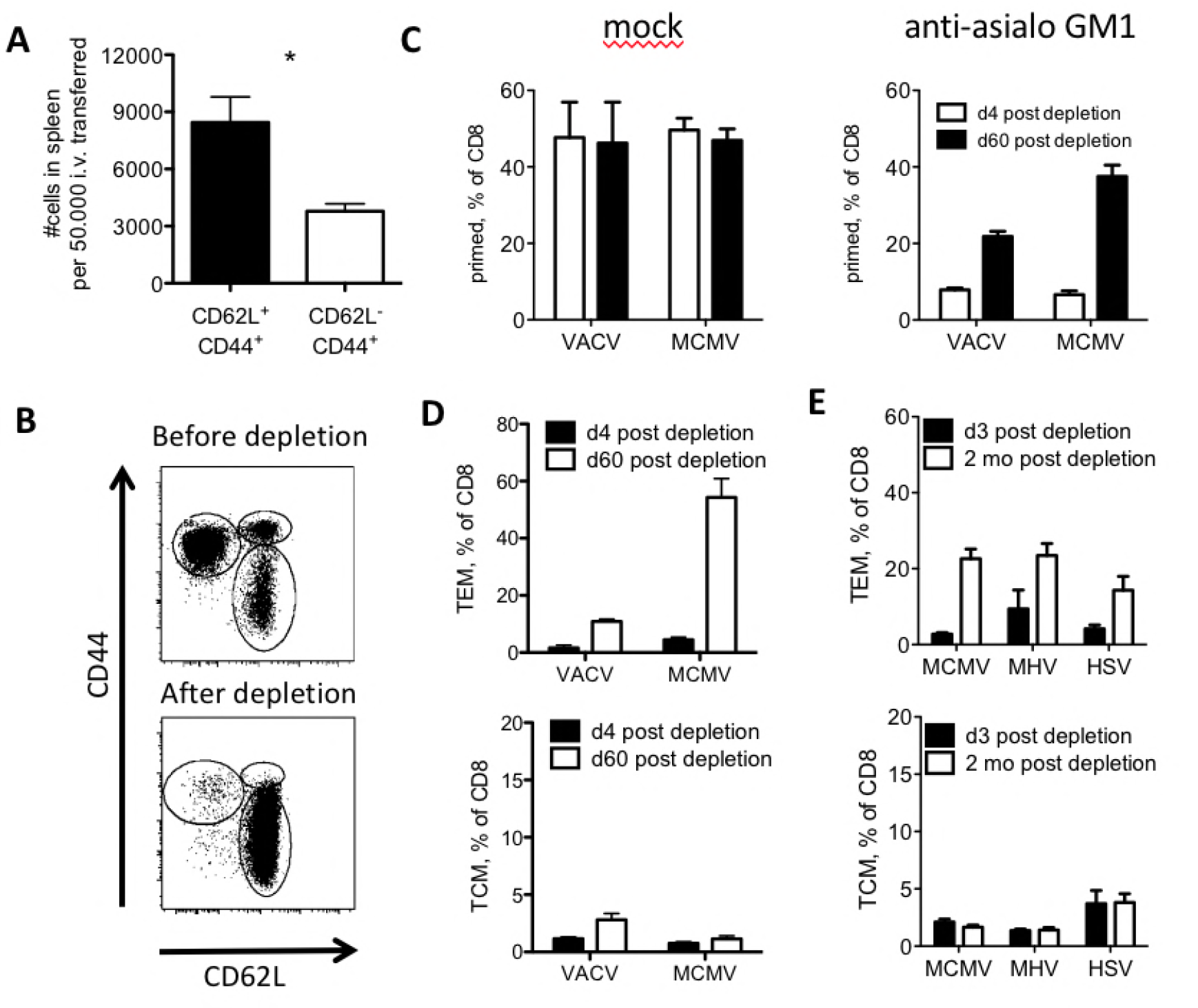
The TEM compartments of herpesvirus-specific CD8 T-cells are restored after depletion with anti-asialo GM1. (A) Ly5.1 C57BL/6 mice were infected with 10^4^ PFU MCMV (Smith strain). At 30 dpi the mice were sacrificed and CD62L^+^ CD44^+^ (TCM) and CD62L^-^ CD44^+^ (TEM) CD8 T-cells were sorted from spleens and transferred into Ly5.2 C57BL/6 recipients (n=4). 18h post transfer the number of Ly5.1+ cells per 50 000 transferred cells was determined in the spleens of the recipients. (B-D) C57CL/6 mice were injected with 10^5^ PFU MCMV-gB, or 10^6^ PFU Vaccinia-gB. 2 months post infection mice were either treated weekly with anti-asialo GM1 antibody or mock treated for 5 consecutive weeks. 4 days and 2 months after the last round of depletion the mice were bled and cells were stained with antibodies against CD8, CD44, CD62L, and CD11a. (B) Representative dot plots of CD8 T-cell subsets before and after depletion. (C) Percentage of primed (CD11a^+^CD44^+^) cells in the CD8 population. (D) Percentages of TEM (CD62L^-^CD44^+^) or TCM (CD62L^-^CD44^+^) cells. 3-5 mice per group. (E) C57CL/6 mice were infected with 2 × 10^5^ PFU MCMV IE2-SSIEFARL, 1 × 10^6^ PFU MHV-68 (MHV) or 2 × 10^5^ PFU HSV-1 (HSV). 10 months post infection the mice were treated weekly with anti-asialo GM1 antibody or an isotype control for four consecutive weeks. 3 days and 2 months after the last round of depletion the mice were bled and cells were stained with antibodies against CD8, CD44, and CD62L. Percentages of TEM and TCM among CD8 T cells were determined. 6 – 10 mice per group; bars indicate means, error bars indicate SEM. Significance was assessed by Mann-Whitney test. *=p<0.05, **=p<0.01, ***=p<0.001, and ****=p<0.0001

### The effector-memory compartment is restored more rapidly than the central-memory after depletion of primed CD8 T-cells with anti-asialo GM1

While all previous experiments suggested ongoing proliferation of effector T-cells in latently infected mice, they did not allow a head-to-head comparison of proliferation in the effector-memory and the central-memory compartment. Adoptive transfer of cell subsets is a method of choice to measure such activity, but it was not suitable due to the homing bias of the transferred subsets. Therefore, to validate if TEM or TCM would replicate more rapidly, we developed an assay where most of the primed CD8 T-cell compartment is depleted upon which we monitored the ability of T-cells to repopulate this compartment. Asialo-GM1 antibodies are widely used for the depletion of NK cells (29, 30), but were shown to also target virus-specific CD8 T-cells (31). We used repeated weekly i. p. injections of asialo-GM1 antibodies starting at 60 dpi and proceeding for four consecutive weeks to deplete the entire CD44^+^CD62L^+^ (TCM) and CD44^+^CD62L^-^ (TEM) compartment, but leave the naïve subset (CD44^-^CD62L^+^) intact (Fig. 4B). Thereupon, we analyzed the fraction of total primed CD8 T-cells at 4 days and 2 months after the last depletion. Antibody administration severely reduced the fraction of primed cells in MCMV and VACV infection, but the cells rebounded at 2 months post depletion, and this was more pronounced in the group infected with MCMV (Fig. 4C). We next defined if the rebound at 2 months was evenly distributed among TCM and TEM cells or more pronounced in any of the subsets, and observed a prominent restoration of TEM cells in MCMV and VACV infection, but a very weak rebound of TCM cells (Fig. 4D). To understand if this effect may be observed in long-term latency and in other latent infections, mice were infected with MCMV, murine herpesvirus clone 68 (MHV-68), or herpes simplex virus type 1 (HSV-1), allowed to establish latency, and treated with asialo-GM1 at 10 months post infection. As in the previous experiment, the depletion strongly reduced the number of primed cells, but their numbers were substantially increased 2 months later. Even more interestingly, the rebound of primed CD8 T-cells was almost exclusively due to the TEM fraction, and essentially no rebound was seen among TCM cells (Fig. 4E). In conclusion, our data showed that in conditions of competition, the TEM cells proliferate more rapidly than TCM cells upon in vivo depletion, and that this is a feature that can be observed in other herpesviral infections as well.

### MCMV-specific CD8 T-cells show antigen-driven proliferation in latently infected mice

To test if the proliferation of effector CD8 T-cells in latently infected mice reflects the proliferation of antigen-specific cells, we analyzed the cycling properties of CD8 T-cells recognizing the well-characterized inflationary M38 antigen. M38-specific CD8 T-cells were identified using flow cytometry via the binding of an M38-MHCI-tetramer, and analyzed using Ki67, FXcycle, and Bcl2 staining (Fig. 5A).

**Figure 5:**
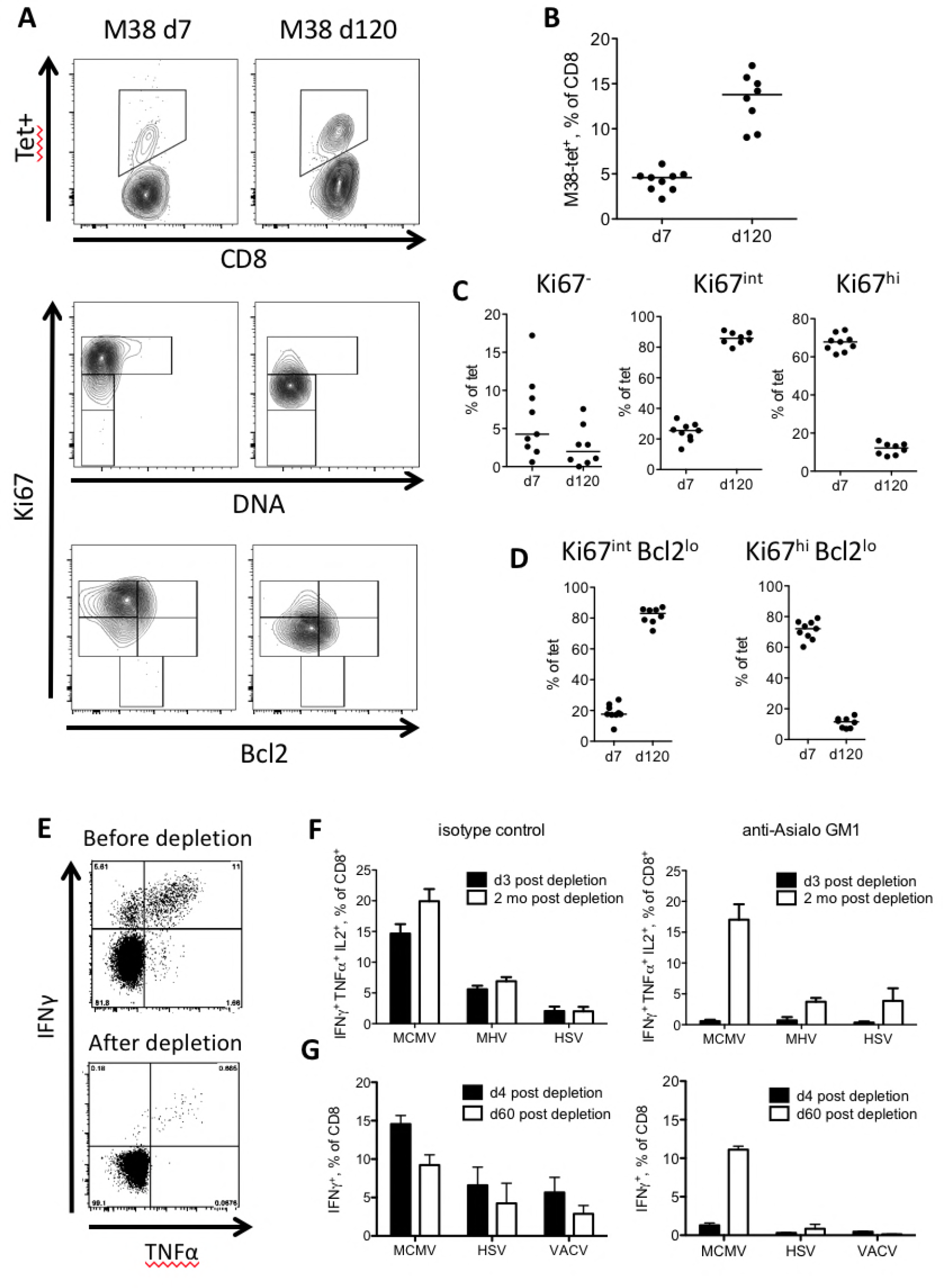
MCMV-specific T-cells show antigen driven proliferation in latently infected mice. 129/Sv mice were infected with 2 × 10^5^ PFU MCMV or VACV. At 7 and 120 dpi, blood was taken and proliferating T-cell subsets were analysed with antibodies directed against CD8, Ki67, and Bcl2. In addition, the DNA content of the cells was determined using the DNA marker FXcycle violet. T-cells specific for M38 were identified via MHC-tetramer staining. (A) Representative dot plots demonstrating specific T-cells among total CD8 T-cells, the gating of cell cycle subsets (Ki67 and FXcycle violet), and proliferating subsets driven by homeostatic or antigen-driven proliferation (Ki67 Bcl2) in tet+ CD8 T-cells. (B) Percentage of tetramer positive CD8 T-cells at 7 and 120 dpi. (C) Percentage of cell cycle subsets among tet^+^ CD8 T-cells. (D) Percentage of Ki67 Bcl2 subsets among tet+ CD8 T-cells. 8-9 mice per group, lines indicate median. (E, F) C57CL/6 mice were infected with 2 × 10^5^ PFU MCMV IE2-SSIEFARL, 1 × 10^6^ PFU MHV-68 (MHV) or 2 × 10^5^ PFU HSV-1 (HSV). 10 months post infection the mice were treated weekly with anti-asialo GM1 antibody or isotype control for four consecutive weeks. 3 days and 2 months after the last round of depletion the mice were bled and cells were stimulated with peptide pools in the case of MCMV (pool of M38 and m139 peptides) and MHV-68 (pool of MHV-68_ORF6_ and MHV-68_ORF6_) or a single peptide (SSIEFARL derived from gB) in the case of HSV-1 and stained with antibodies against CD8, IL-2, IFNγ, and TNFα to identify cytokine producing CD8 T-cells. (E) Representative dot plots of cytokine-producing MCMV specific CD8 T-cells before and after depletion with anti-asialo GM1 antibody. (F) Percentages of cytokine producing CD8 T-cells in isotype and anti-asialo GM1 treated mice. 6-10 mice per group. (G) C57CL/6 mice were infected with 2 × 10^5^ PFU MCMV-ie2-SSIEFARL, 2 × 10^5^ PFU HSV-1 or 2 × 10^5^ PFU Vaccinia-SSIEFARL. 2 months post infection the mice were treated weekly with anti-asialo GM1 antibody or isotype control for four consecutive weeks. 4 days and 60 days post depletion the mice were bled and cells stimulated with the HSV-1 glycoprotein-derived epitope gB (SSIEFARL) and stained with anti-CD8 and anti-IFNγ. The CD8 T-cells were analysed for IFNγ expression. 6-10 mice per group, bars indicate means, error bars indicate SEM.

Consistent with prior reports (32), M38-specific cells expanded from 4.6% of the CD8 T-cell pool at 7 dpi to 14% at 120 dpi (Fig. 5B). Very few M38-specific cells were low in Ki67 expression both at 7 and at 120 dpi. The percentage of Ki67^int^ T-cells increased from a mean of 26% at 7 dpi to 86% at 120 dpi, while the percentage of Ki67^hi^ T-cells decreased from 68% on average at 7 dpi to 12% at 120 dpi (Fig. 5C). Similarly, M38-specific CD8 T-cells were mainly Ki67^hi^Bcl2^lo^ (72%) at 7 dpi, whereas at 120 dpi most T-cells were Ki67^int^Bcl2^lo^ (83%). 12% retained the Ki67^hi^ Bcl2^lo^ phenotype (Fig. 5D). Thus, while the data resembled the time-associated pattern that was observed in the total CD8 T-cell pool, a larger fraction of M38-specific cells were Ki67^hi^ at 120 dpi, than the data observed in the total CD8 T-cell pool (median of 6.2%, Fig. 1B). The immunodominant peptide (RALEYKNL) derived from the ie3 MCMV gene (33) showed similar phenotypes (data not shown). Taken together, the results suggest that CD8 T-cells recognizing immunodominant inflationary antigens may vigorously proliferate during MCMV latency.

### Virus-specific CD8 T-cells are restored after asialo-GM1 depletion

To investigate if virus-specific inflationary CD8 T-cells would rebound after the antibody mediated depletion of primed cells, blood lymphocytes from mice latently infected with the three herpesviruses (MCMV, MHC-68, HSV-1; see Fig. 4E) were restimulated with viral peptides, and the responding cells were quantified by intracellular cytokine staining and flow cytometry. We used peptide pools in the case of MCMV (pool of M38 and m139 peptides) and MHV-68 (pool of MHV-68_ORF6_ and MHV-68_ORF6_) or a single peptide (SSIEFARL derived from gB) in the case of HSV-1. Asialo-GM1 treatment resulted in highly efficient depletion of antigen-specific cells (Fig. 5E), yet the fraction of virus-specific CD8 T-cells was nearly completely replenished two months later. This was observed in MCMV, MHV-68 and HSV-1 infected mice (Fig. 5F), arguing that inflationary CD8 T-cells in herpesvirus infections are capable of replenishing their pool after depletion.

To understand if the restoration of virus-specific CD8 T-cells is typical for persistent infections, or whether it also occurs in non-persistent infection, we compared mice infected with MCMV and HSV-1 to mice infected with VACV. We normalized the populations of CD8 T-cells responding to antigen by using recombinant MCMV and VACV clones expressing the HSV-1 epitope SSIEFARL (34, 35). This enabled us to compare specific T-cell responses in persistent herpesvirus infections and a non-persistent infection, while excluding the possibility that peptide-intrinsic differences, such as differences in avidity of responding CD8 populations, may influence the outcome. We treated the mice with asialo-GM1 antibodies at 60 dpi and analyzed SSIEFARL-specific CD8 T-cell responses at 4 days post depletion or two months later. The depletion of virus-specific CD8 T-cells was highly efficient in all three virus infections, but their numbers were restored only in MCMV, and partly in HSV-1 infection, whereas VACV-infected mice showed essentially no recovery (Fig. 5G). Taken together, the results confirmed that the proliferative potential of CD8 T-cells in persistent infections is likely due to antigen-driven proliferation that cannot be observed in infections that are cleared.

It is important to note that adoptive transfer of M38- or m139-specific CD8 T-cells sorted for the CD62L^+^ or CD62L^-^ phenotype from spleens of latently infected Ly5.1 C57BL/6J mice into naïve Ly5.2 C57BL/6J mice also showed significantly more homing of CD62L^+^ CD8 T-cells to the spleen compared to CD62L^-^ CD8 T-cells (Supplementary Fig. 3). Hence, we avoided testing the proliferative capacity of antigen-specific cells by adoptive transfer.

### Dynamic monitoring reveals robust cycling of inflationary CD8 T-cells

The comparison of Tam treated and mock-treated R26 CreER^T2^ mice revealed significantly smaller fractions of Ki67^hi^ subsets in Tam treated mice, especially in the effector compartment of CD8 T-cells (Fig. 3A, B, C). To define in longitudinal experiments the effect of Tamoxifen treatment on the inflationary responses, we analyzed the kinetics of CD8 T-cell responses in the blood of MCMV infected R26 CreER^T2^ mice before and after Tamoxifen treatment, and compared it to mock-infected controls. Mice were bled at 7, 14, 28, 60, 90 and 120 days post MCMV infection, before Tam containing food was administered for 28 days. Subsequently, mice were bled at 150, 180 and 210 dpi. Blood lymphocytes were restimulated with the inflationary peptide RALEYKNL from the MCMV gene ie3 (32) and analyzed for surface marker and intracellular cytokine expression by flow cytometry. First, we analyzed the effector cells (CD127^-^CD44^+^) and observed a stable population of the CD8 compartment until 120 dpi, followed by a marked drop after 28 days of Tamoxifen treatment. This was not observed in the untreated control group (Fig. 6A). The reduction of the ie3-specific CD8 T-cell fraction was even more pronounced (Fig. 6B). Once the Tam-spiked food pellets were replaced with standard diet, the analyzed CD8 subsets rebounded (Fig. 6A, B). Taken together, the data confirmed that Tam treatment reduced the effector subset of T-cells in general and antigen-specific inflationary cells in particular, arguing strongly for continuous cycling of these subsets. Tam administration in latently infected C57BL/6 mice resulted in no loss of TEM or antigen-specific CD8 T-cells (Supplementary Fig. 4A-D), arguing that the effects were specific for Tam effects in R26 CreER^T2^ mice. In an independent experiment, we administered Tamoxifen at 60 dpi to mice infected with the SSIEFARL-expressing MCMV (see Fig. 5F), and we observed a reduction of T-cells specific for the inflationary SSIEFARL and the M38 epitope (Supplementary Fig. 5A, B).

**Figure 6:**
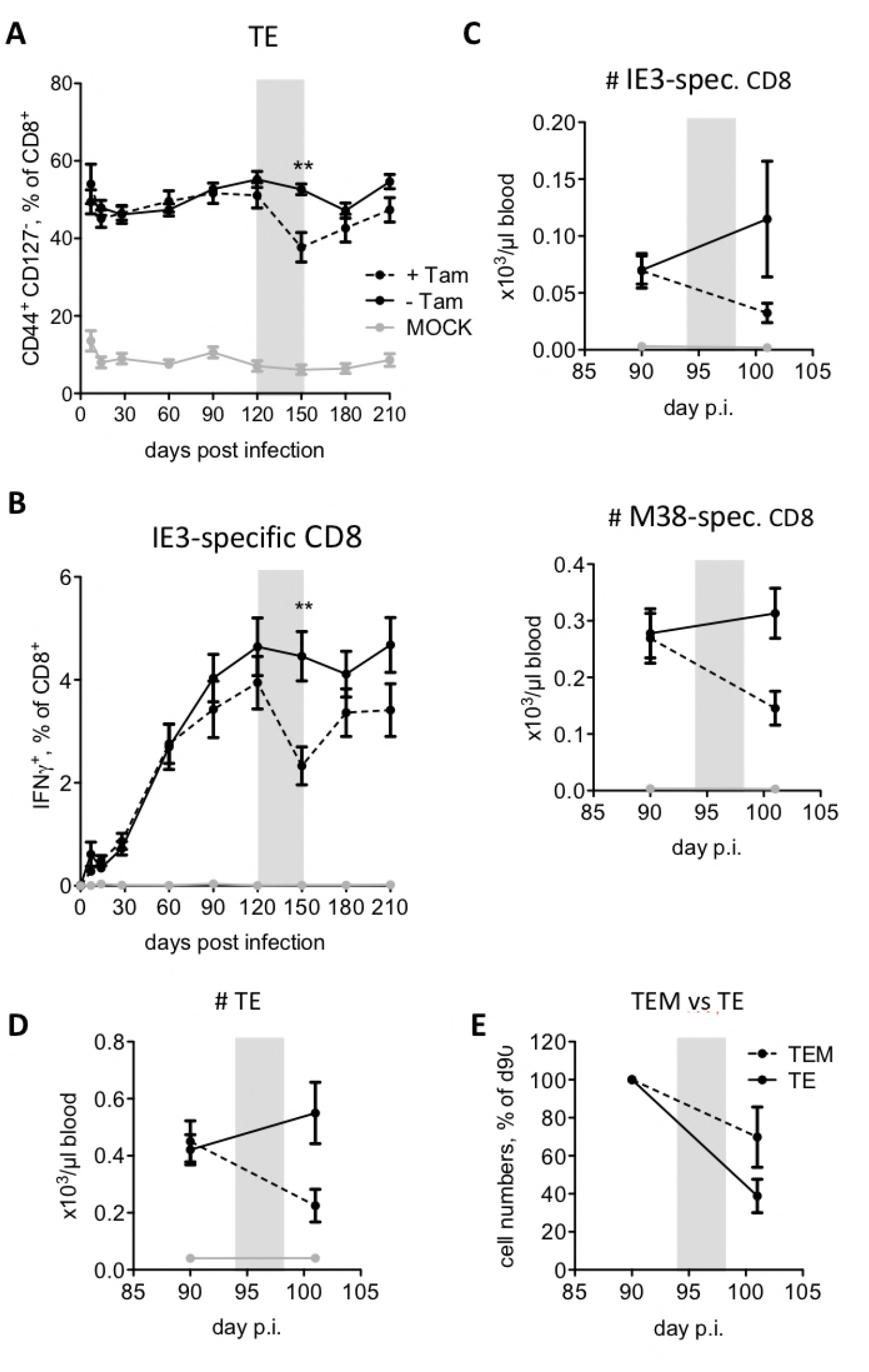
MCMV-specific CD8 T-cells are depleted by Tamoxifen treatment in R26 CreER^T2^ mice. (A, B) R26 CreER^T2^ mice were infected with 10^6^ PFU MCMV or 200 μl PBS (mock). 4 months post infection half of the mice received food pellets containing 400 mg/kg Tamoxifen for 4 weeks (indicated with a grey rectangle). Blood was collected and analysed at 0, 7, 14, 28, 60, 90, 120, 150, 180, and 270 dpi. Blood leukocytes were stimulated with the IE3 peptide (RALEYKNL) and stained for CD8, CD44, CD127, and IFNγ expression. (A) Kinetics of effector (CD44^+^ CD127^-^, TE) CD8 T-cells before, during, and after Tamoxifen treatment. (B) Percentages of IFNγ^+^ CD8 T-cells specific for the IE3 peptide before, during, and after Tamoxifen treatment. Data are pooled from three independent experiments with up to 15 mice per group in total. Error bars indicate SEM. Significance was assessed by Mann-Whitney test. \*\**p* < 0.01. \*\*\*\**p* < 0.0001. (C, D) R26 CreER^T2^ mice were infected with 10^6^ PFU MCMV or 200 μl PBS (mock). 3 months post infection mice half of the mice were treated with 80mg/kg bodyweight Tamoxifen by oral gavage on five consecutive days (d94-d98). Blood was collected and analysed on d90 (pre treatment) and on day 101 (3 days post Tamoxifen). Lymphocyte counts in blood were acquired via Vetscan. (C) Numbers of M38 and IE3-specific T-cells were determined by tetramer-staining. (D) Counts of effector (CD44^+^ CD127^-^, TE) CD8 T-cells. Data from 2 independent experiments were pooled, with up to 15 mice per group in total. Dots indicate means, error bars indicate SEM. (E) Comparison of TEM and TE loss. Cell counts in each subset prior to Tamoxifen were normalized as 100% and post-depletion numbers are shown. 2 experiments with 15 mice per group were pooled. Data are means, error bars and SEM

Finally, we considered the possibility that the percentagewise loss of effector cells could have been a reflection of absolute gains of cells in other subsets of T-cells, rather than a loss of antigen-specific cells with effector phenotypes. While unlikely, this scenario could have mimicked as a relative loss of effector and antigen-specific T-cells. To exclude this possibility, we repeated the kinetics experiment and measured the relative and absolute counts of CD127^-^CD44^+^ CD8 T-cells and of CD8 T-cells specific for two inflationary epitopes derived from the ie3 and the M38 epitope. As in the previous experiment, Tam feeding decreased the relative fraction of antigen-specific and effector cells (data not shown), and this was reflected in a decrease of absolute counts of antigen-specific CD8 T-cells (Fig. 6C), as well as effector CD8 T cells (Fig. 6D). Importantly, the drop in absolute counts of TE cells was stronger than the loss of the TEM subset (Fig. 6E), consistent with the high percentage of Ki67^hi^ cells (Fig. 2B) and cycling within the TE cells. In conclusion, selective targeting of cycling cells for elimination resulted in a selective loss of cells expressing high levels of Ki67, a reduction of antigen-specific T-cells recognizing inflationary epitopes and of the effector CD8 T-cell fraction in general.

## Discussion

In our study, we document a robust and long-term proliferation of effector T-cells in latent herpesviral infection. In light of the previously published literature, which classified this subset as short-lived effector cells that are maintained by continuous influx from naïve and central-memory cells, our results are surprising. Two groups have independently shown that MCMV-specific TEM T-cells poorly proliferate upon transfer into MCMV-infected mice (19, 20), whereas recent adoptive transfer data by the Robey lab argued that memory (KLRG1^-^) or intermediate (KLRG1^+^) CXCR3^+^ cells maintain the effector compartment in chronic Toxoplasma infection (36). We show that effector cells robustly cycle and that their own cycling substantially contributes to their maintenance in chronic infection. We explain this stark difference in conclusions by differences in the experimental approach. Namely, we show that adoptive transfer experiments introduce a bottleneck for T-cell expansion. This bottleneck depends on the ability of transferred cells to home to secondary lymphoid organs, resulting in an inherent – and previously unappreciated – bias towards the expansion of cells expressing defined chemokine receptors or integrins, such as CD62L. We do not challenge the conclusion that TCM proliferate better than TEM upon adoptive transfer, and that immunotherapy by adoptive transfer of TCM cell preparations is likely to yield better clinical outcomes (37). However, we show that adoptive transfer may bias for cell survival and homing during the transfer, rather than merely reflecting cell proliferation after the transfer. Considering the widespread use of this method, our data invites caution in extrapolating adoptive transfer results to conditions of cell proliferation in their natural niche. To exclude that our main conclusion is likewise influenced by a bias in the method, we performed four independent approaches, based on (i) precise Ki67 staining, (ii) in vivo deuterium labeling, (iii) targeted depletion of TCM and TEM, and (iv) targeted depletion of cycling cells. All four methods pointed to robust proliferation of effector CD8 T cells and antigen-specific CD8 T cells in chronically infected mice, especially in the spleen. Therefore, we conclude that effector cell cycling substantially contributes to the maintenance of inflationary responses in steady state conditions.

We observed a stark but underreported three-way separation in Ki67 expression patterns in lymphocytes. The vast majority of published literature does not account for the large Ki67^int^ subset, which overlaps with the primed CD8 T-cell compartment. We assume that we obtained an optimal cell separation by using a cell permeabilization procedure tailored for the detection of transcription factors, which allowed us to accurately stain the intra-nuclear Ki67. CreER^T2^-mediated depletion did not affect the size of the Ki67^int^ subset, and their genomes were diploid, arguing that these cells are largely quiescent, while cycling is specific for the Ki67^hi^ subset. This is in line with a recent publication that reported somewhat diminished Ki67 levels in CD4 T-cells that had previously cycled, but were not in cell cycle at the time of analysis (38). Consequently, conclusions from numerous previous studies (including our own ones) may require revisiting in light of this optimized labeling.

We base our conclusions on several complementary methods that pointed us in the same direction. We used *in vivo* deuterium labeling to measure cell proliferation and turnover and observed that the TEM subset was the fastest one on both accounts. Antibodies recognizing asialo-GM1 are a mainstay of NK-cell depletion in experimental mouse models, but they also have the potential to target and deplete T-cells specific for viral antigens as shown by others previously (31). We showed here that essentially all primed CD8 T-cells express asialo-GM1; thus, they may be depleted by rigorous and repeated antibody administration, yet the TEM cells are the first one to recover from this depletion. Similarly, we are not the first ones to report that loxP-like sequences present in host genomes may be recognized by the Cre-recombinase (26) resulting in the depletion of rapidly cycling hematopoietic cells (27). However, we are to our knowledge the first ones to exploit this insight to deplete specifically the antigen-specific cells that proliferate and thus define which populations are maintained by renewal and which ones are long-term quiescent. Taken together, our results indicate robust cycling in the effector CD8 T-cell subset of mice latently infected with herpesviruses. Considering that latent herpesvirus infections are ubiquitous and found in mammals and birds, it is reasonable to assume that in a natural microbiological environment such proliferation occurs in all mammals, including all people worldwide. Therefore, our results appear to bear profound implications for our understanding of T-cell population dynamics in natural settings.

## Material and Methods

### Mouse strains

129S2/SvPas Crl (129/Sv) mice were purchased from Charles River (Sulzfeld, Germany). C57BL/6 mice were purchased from Janvier (Le Genest Saint Isle, France). Rosa26-CreER^T2^ (R26CreER^T2^) mice on the C57BL/6 background were bred at the Helmholtz Centre for Infection Research (Braunschweig, Germany). Ly5.1 and Ly5.2 C57BL/6 mice were maintained at the Central Animal Facility of Leiden University Medical Center. Mice were 8-10 weeks old at the beginning of an experiment, R26CreER^T2^ mice were used at 8-20 weeks of age. Mice were housed and handled in accordance with good animal practice as defined by Federation of Laboratory Animal Science Associations and the national animal welfare body Die Gesellschaft für Versuchstierkunde/Society of Laboratory Animals.

### Ethics statement

All animal experiments were approved by the responsible state office (Lower Saxony State Offi'ce of Consumer Protection and Food Safety, Germany; permit no. 33.9-42502-04-10/0109, 33.9-42502-04-11/0426 and 33.9-42502-04-15/1832) or by the Animal Experiments Committee of LUMC (Leiden University Medical Center, The Netherlands; DEC 12070). Animal care and use protocols adhered to Directive 2010/63/EU.

### Cells

M2-10B4 (CRL-1972), Vero (CCL-81), and NIH 3T3 fibroblasts (CRL-1658) (all from American Type Culture Collection) were maintained in DMEM supplemented with 10% FCS, 1% glutamine, and 1% penicillin/streptomycin. C57BL/6 murine embryonic fibroblasts (MEFs) were prepared in our lab from 14 days pregnant mice and maintained as described previously (39).

### Viruses

BAC-derived wild-type MCMV (MCMV WT) and MCMV-SSIEFARL were propagated as described previously (40) and the virus stocks were prepared from M2-10B4 lysates purified on a sucrose cushion as described previously (41). Virus titers were determined on MEFs by plaque assay. HSV-1 strain 17 obtained from Dr. J. Nikolich-Zugich (University of Arizona, Tucson, AZ) was grown and titrated on Vero cells (42). The MHV-68 was kindly provided by Dr. H. Adler (43), Helmholtz Zentrum München, and the virus stock was produced and titrated on NIH3T3 cells. Western Reserve Vaccinia virus (VACV) and VACV-SSIEFARL were grown and titrated on VERO cells (44). Mice were infected with the indicated dose of purified, tissue culture-derived virus and housed in specific pathogen-free conditions throughout the experiment.

### Cell surface staining

Blood cells were surface stained for 30 min at 4°C with the following antibodies, depending on the experiment: anti-CD4-Pacific Blue (clone GK1.5; BioLegend), anti-CD4-allophycocyanin (clone GK1.5; BioLegend), anti-CD4-PE-Cy7 (clone GK1.5; BioLegend), anti-CD8a-PerCP/Cy5.5 (clone 53-6.7; BioLegend), anti-CD44-Alexa Fluor 700 (clone IM7; BioLegend), anti-CD62L-eFluor 605NC (clone MEL-14; eBioscience), anti-CD127-PE (clone A7R34; Biolegend), anti-CD62L-Brilliant Violet 510 (clone MEL-14; Biolegend), anti-CD127-PE-Cy7 (clone A7R34; Biolegend), and anti-CD3-allophycocyanin-eFluor 780 (clone 17A2; eBioscience). Antigen-specific cells were detected with allophycocyanin-labeled MHCI-peptide tetramers specific for M38, m139, or gB (see also Peptides) or oligomerized streptamers specific for IE3. Briefly, the MHC class I-peptide-Strep molecules were first oligomerized (incubation for 45 min at 4°C with Strep-Tactin allophycocyanin; IBA Life Sciences) before cells were stained with the oligomerized MHC class I-Strep molecules for 15 min at 4°C with subsequent staining with surface antibodies for additional 30 min.

### Ki67, Bcl-2, and FxCycle staining

For intracellular staining of Ki67 and Bcl-2, cells were first surface stained and then fixed for 20 min at room temperature with 100μl fixation/permeabilization buffer of the FoxP3/Transcription factor staining set (eBioscience) followed by 15 min at room temperature in 100μl permeabilization buffer (eBioscience). Subsequently the cells were stained with Ki67-PE (clone 16A8, Biolegend) and Bcl-2-AF488 (clone BCL/10C4, Biolegend) in 100μl permeabilization buffer for 30 min at room temperature. After the intracellular staining cells were resuspended in 100μl of FXcycle staining solution (FxCycle Violet stain, life technologies) and incubated for 30 min at room temperature before acquisition with an LSR Fortessa (BD Biosciences). Cytometric results were analyzed with FlowJo software (version 9.8.3).

### AnnexinV and 7AAD staining

4 x 10^6^ splenocytes or 300μl blood were used for the apoptosis staining. The cells were stained first with a Live/Dead marker (LIVE/DEAD^®^ Fixable Blue Dead Cell Stain Kit, Invitrogen) for 30 min at 4°C, followed by a washing step and then stained with CD8-eFluor 450 (clone 53-6.7; eBioscience), CD62L-PE (clone MEL-14; BD) and CD44-FITC (clone IM7; BD) for 15 min at 4°C. After an additional washing step, the cells were re-suspended in 250 μl binding buffer, before 5 μl Annexin V-APC (BD Biosciences) and 2.5 μl 7AAD (Sigma-Aldrich) were added and incubated for 15 min at RT. 250 μl binding buffer was added to the staining mix and the cells were acquired in an LSR-II or an LSR Fortessa cytometer (BD Biosciences). Cytometric results were analyzed with FlowJo software (version 9.5.3).

### Intracellular cytokine staining

For intracellular cytokine staining, cells were first surface stained and then subsequently fixed for 5 min with 100 μl IC fixation buffer (eBioscience), followed by 3 min permeabilisation with 100 μl permeabilisation buffer (eBioscience) and overnight incubation with anti-IFN-γ-allophycocyanin (clone XMG1.2; BioLegend), anti-TNF-α-FITC (clone MP6-XT22; Biolegend) and anti-IL-2-Pacific Blue (clone JES6-5H4; BioLegend). Cells were washed and acquired in an LSR-II cytometer (BD Biosciences). Cytometric results were analyzed with FlowJo software (version 9.5.3).

### Stable isotope labeling

120 days after infection, 25-week old 129/Sv mice received 8% deuterated water (99.8% D_2_O, Cambridge Isotope Laboratories) in their drinking water for 28 days. At day 7, mice were given a boost injection (i.p.) of 15ml/kg D_2_O in phosphate-buffered saline (PBS).

### Cell preparation and sorting

Spleens, thymocytes, and blood were isolated at different time points during and after label administration. Blood was collected in a vial containing anti-coagulant EDTA and spun down to isolate plasma. Plasma was stored at −80°C until analysis. Organs were mechanically disrupted (using a plunger and a cell strainer) to obtain single-cell suspensions. Splenocytes were stained with monoclonal antibodies: anti-CD3e-APC-FITC (clone 145-2C11; BD-Biosciences), anti-CD4-pacific blue (clone GK1.5; BioLegend), anti-CD8a-Percp/cy5.5 (clone 53-6.7; BioLegend), anti-CD44-Alexa fluor 700 (clone IM7; BioLegend), and anti-CD62L-evolve605 (clone MEL-14; eBioscience). Within CD8^+^ tetramer-negative splenocytes, naïve T-cells were defined as CD62L^+^CD44^-^, central-memory T-cells as CD62L^+^CD44^+^, and effector-memory T-cells as CD62L^-^CD44^+^. Cells were sorted using an Aria-II SORP or MoFlo XDP cell sorter. Genomic DNA was isolated from thymocytes of age-matched, uninfected mice and sorted T-cell subsets from MCMV-infected mice according to the manufacturer’s instructions using the NucleoSpin Blood QuickPure kit (MACHEREY-NAGEL), and stored at −20°C until further analysis.

### Measurement of deuterium enrichment in DNA and body water

Deuterium enrichment in plasma (as a measure of deuterium availability in the body water) of uninfected mice and in the DNA of the different cell subsets was analyzed by gas-chromatography/mass-spectrometry (GC/MS) using an Agilent 5973/6890 GC/MS system (Agilent Technologies). Deuterium enrichment in DNA was measured according to (4). Briefly, DNA was hydrolyzed to deoxyribonucleotides and derivatized to penta-fluoro-triacetate (PFTA). The derivative was injected into the GC/MS equipped with a DB-17 column (Agilent Technologies) and measured in SIM mode, monitoring ions m/z 435 (M+0), and m/z 436 (M+1). From the ratio of ions, deuterium enrichment in the DNA was calculated by calibration against deoxyadenosine standards of known enrichment. Plasma was derivatized to acetylene (C_2_H_2_) as previously described (4). The derivative was injected into the GC/MS equipped with a PoraPLOT Q 25 × 0.32 column (Varian), and measured in SIM mode, monitoring ions m/z 26 (M+0) and m/z 27 (M+1). From the ratio of ions, deuterium enrichment in the plasma was calculated by calibration against standard water samples of known enrichment.

### Adoptive T-cell transfer

For the adoptive T-cell transfer, CD62L^-^ and CD62L^+^ CD44^+^ CD8^+^ T-cells from spleens of latently infected Ly5.1 mice were sorted after CD8 enrichment. 50.000 CD62L^+^ or 50.000 CD62L^-^ cells were transferred into naïve Ly5.2 mice. 3h after transfer, the number of transferred cells present in the spleen were determined by flow cytometry.

### Depletion of memory T-cells

Mice were i.p. injected with 100 μg purified anti-Asialo GM1 (polyclonal; eBioscience) or 100 μg rabbit anti-human IgG (polyclonal; Thermo Scientific) weekly for 4 weeks and housed in specific pathogen-free conditions throughout the experiment.

### Tamoxifen application

Tamoxifen (Tamoxifen, T5648-1g, Sigma) was dissolved in an Ethanol/corn oil (C8267-500ML, Sigma) mixture (1/49) at a concentration of 20mg/ml. Mice were weighed and received 80mg/g BW Tamoxifen via orally on 5 consecutive days, followed by 3 days break, and 1 more day of treatment. Control animals received corn oil only. For long-term treatment, Tamoxifen was administrated orally with pellets containing 400 mg/kg Tam CRD Tam^400^/CreER (Teklad) at indicated time points.

### Peptides

The peptides m139 (H-2K^b^-restricted, ^419^TVYGFCLL^426^), IE3 (H-2K^b^-restricted, ^416^RALEYKNL^423^), M38 (H-2K^b^-restricted, ^316^SSPPMFRV^323^) (11), the HSV-1 glycoprotein-derived epitope gB (H-2K^b^-restricted, ^498^SSIEFARL^505^) (45), the MHV-68_ORF6_ (H2-D^b^-restricted, ^487^AGPHNDMEI^495^) and MHV-68_ORF61_ (H2-K^b^-restricted, ^524^TSINFVKI^531^) were synthesized and HPLC purified (65-95% purity) at the HZI peptide-synthesis platform.

### Peptide stimulation

Cells were stimulated with peptides (1 μg/ml) in 100 μl RPMI 1640 (10% FCS supplemented) for 6 h at 37° C with brefeldin A (Cell Signaling Technology) added at a concentration 10 μg/ml for the last 5 h stimulation. Negative control samples were generated for all tested groups by incubating cells in the same conditions, but in the absence of any peptide. Cells were tested for cytokine responses by intracellular cytokine staining using flow cytometry. Cytokine responses observed in unstimulated samples were considered background noise due to unspecific Ab binding and were subtracted from the values observed in test samples.

### Statistics

Comparisons between two groups were performed using the Mann-Whitney U test (two-tailed). Statistical analysis was performed using the GraphPad Prism program.

### Mathematical modeling of deuterium labeling data

We used a mathematical model to deduce the dynamics of naïve and memory T-cells from the deuterium labeling data. To control for changing levels of deuterium (^2^H) in body water over the course of the experiment, a simple label enrichment/decay curve was fitted to ^2^H enrichment in plasma:

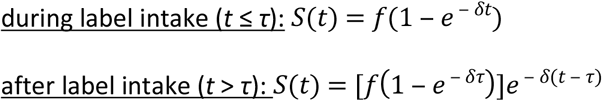

as described previously (46), where *S(t)* represents the fraction of D_2_O in plasma at time *t* (in days), *f* is the fraction of D_2_O in the drinking water, labelling was stopped at *t* = τ days, and *δ* represents the turnover rate of body water per day. The best fit for *S(t)* was used in the labelling equations for the different cell populations (see below). Up- and down-labeling of the thymocyte population was analyzed as previously described (47), to estimate the maximum level of label intake that cells could possibly attain. The best fits to the plasma and thymocyte data are shown in Supplementary Figure 1.

To model the deuterium enrichment in the different T-cell subsets, we used a previously published mathematical model, which in principle allows for kinetic heterogeneity between cells within a population (47). Based on previous observations, we know there is a time delay between T-cell production in the thymus and the appearance of labeled DNA in naïve T-cells in the spleen (47), which we fixed at 4 days. For all subpopulations, we found that the data could be well described with a kinetically homogeneous model, i.e. where all cells have the same average turnover rate *p*, from which we derived the average lifespan as 1/p.

Best fits were determined by minimizing the sum of squared residuals using the R function nlminb, after transforming the fraction of labeled DNA (x) to arcsin(sqrt(x)). The 95% confidence intervals were determined using a bootstrap method where the residuals to the optimal fit were resampled 500 times.

## Acknowledgement

We thank Ilona Bretag, Ayse Barut, and Inge Hollatz-Rangosch for excellent technical assistance and Dirk Busch for providing us streptamer reagents. This work was supported by the ERC-StG 260934, the DFG grant SFB900 TP B2, Helmholtz Association grants VH-NG-638 and VH-VI-424 to LCS, and the European Union Seventh Framework Programme (FP7/2007-2013) under grant agreement 317040 (QuanTI) to JAMB. The funding agencies had no role in the study design, nor in the data analysis and manuscript preparation.

---

**Supplementary Figure 1:**
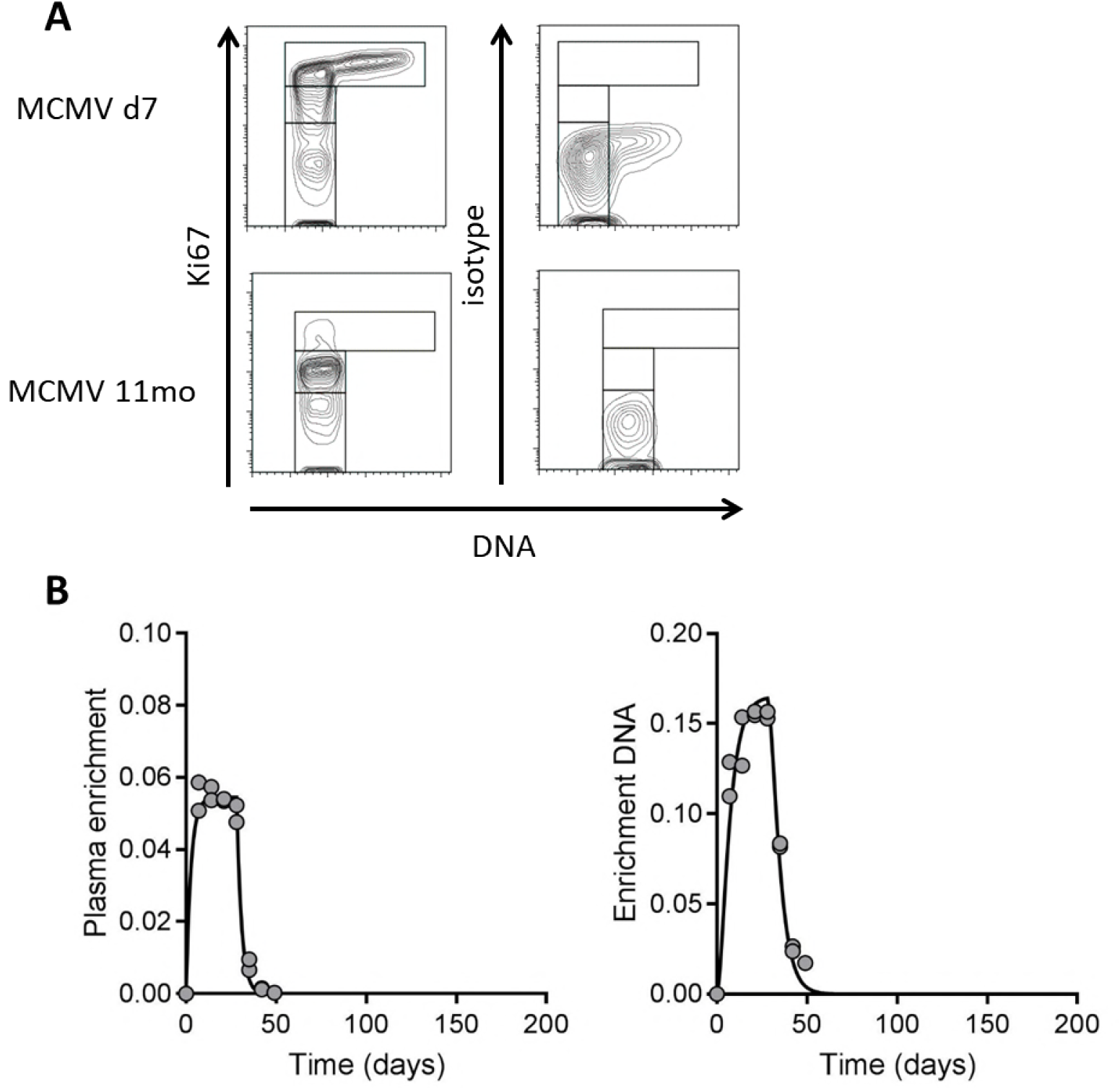
Ki67 staining isotype control and supplemental Deuterium enrichment data. (A) Blood from mice that had been infected with MCMV for either 7 days or 11 months was stained with antibodies against CD3, CD8, and Ki67. Additionally, samples were stained using an isotype control for Ki67 instead. Furthermore, the DNA dye FXcycle violet was added. Representative dotplots are shown. (B) The fraction deuterium labeled plasma (as a measure of deuterium availability in the body water) over time during 4 weeks of D_2_O administration and the subsequent 16 weeks after label cessation (left). The fraction of deuterium labeled DNA in thymocytes over tim (right). Thymocytes were used as a population of rapidly turning over cells to estimate the maximum level of label incorporation that cells could possibly attain. Circles represent measurements from individual mice; curves represent the best fits of the mathematical model to these data (see Methods).

**Supplementary Figure 2:**
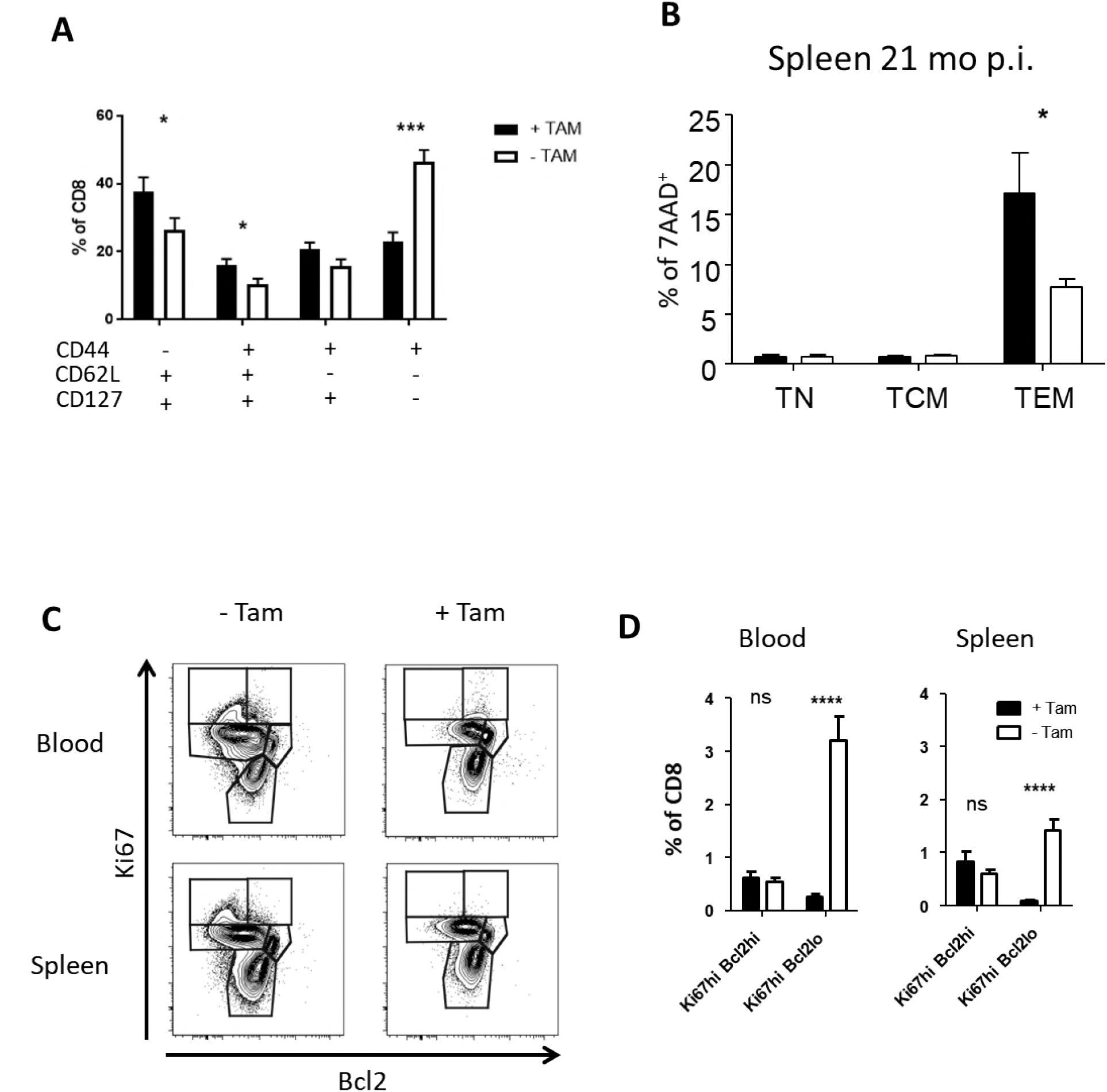
Supplemental Tamoxifen data. Fraction of CD8 subsets in the CD8 pool upon Tam treatment. Mice were MCMV infected and Tam treated as shown in figure 3. Blood leukocytes were stained for CD8 and the indicated surface markers and gated in the CD44-CD62L+CD127+ (naïve) CD44+CD62L+CD127+ (central memory), CD44+CD62L-CD127+ (effector memory) and CD44+CD62L-CD127-(Effector) subsets. Shown are group means + standard deviations from two pooled experiments. (B) Tamoxifen induces cell death in splenic TEM subsets after a life of latency. R26 CreER^T2^ mice were infected with 10^6^ PFU MCMV. 21 months post infection the mice received food pellets containing 400 mg/kg Tamoxifen for 2 weeks. Splenocytes were stained with antibodies against CD8, CD44, CD62L, and 7AAD. Percentage of 7AAD+ cells in immune phenotype subsets are shown. 4 mice per group; significance was assessed by Mann-Whitney test. * = p<0.05. (C+D) Tam treatment depletes primarily Bcl2^lo^ cells. R26CreER^T2^ mice were infected with 10^6^ PFU MCMV. At 94 dpi, the mice received 80mg/kg BW Tamoxifen by oral gavage for 5 consecutive days, followed by one day break, and one more day of treatment, before mice were sacrificed. Cell proliferation in CD8 T-cell subsets was characterized by CD8, Ki67, and Bcl2 expression. (C) Representative dot plots for CD8 T-cell proliferation in blood and splenocytes in treated and untreated mice. (D) Percentages of cell cycle subsets Ki67^hi^ Bcl2^hi^ and Ki67^hi^ Bcl2^lo^ in total CD8 T-cells. Pooled data from two independent experiments (12 mice in total) are shown. Bars indicate means, error bars indicate SEM. Significant differences were determined by Mann-Whitney test, ****=p<0.0001.

**Supplementary Figure 3:**
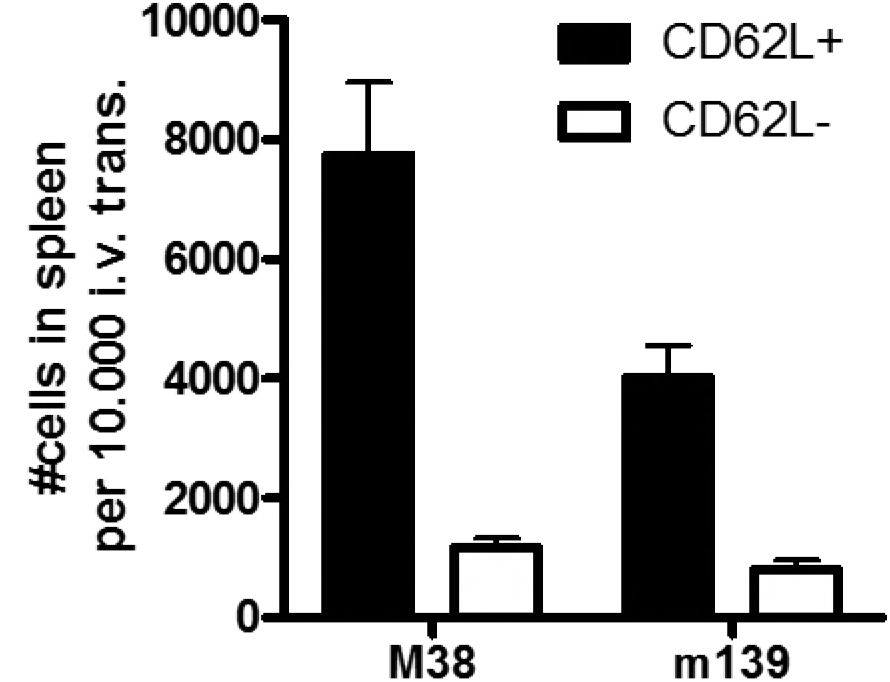
Antigen-specific CD62L^+^ T-cells home to the spleen more efficiently than CD62L^-^. Ly5.1 C57BL/6 mice were infected with 10^4^ PFU MCMV (Smith strain). At 30 dpi the mice were sacrificed and CD62L^+^ tet^+^ and CD62L^-^ tet^+^ CD8 T-cells were sorted from spleens after CD8 T cell enrichment, using tetramers for the inflationary epitopes m139 and M38. Cells were transferred into Ly5.2 C57BL/6 recipients (n=2 for CD62L^+^ tet^+^ and n=4 for CD62L^-^ tet^+^). 18h post transfer the number of Ly5.1+ cells per 10 000 transferred cells was determined in the spleens of the recipients. Bars indicate means, error bars indicate SEM.

**Supplementary Figure 4:**
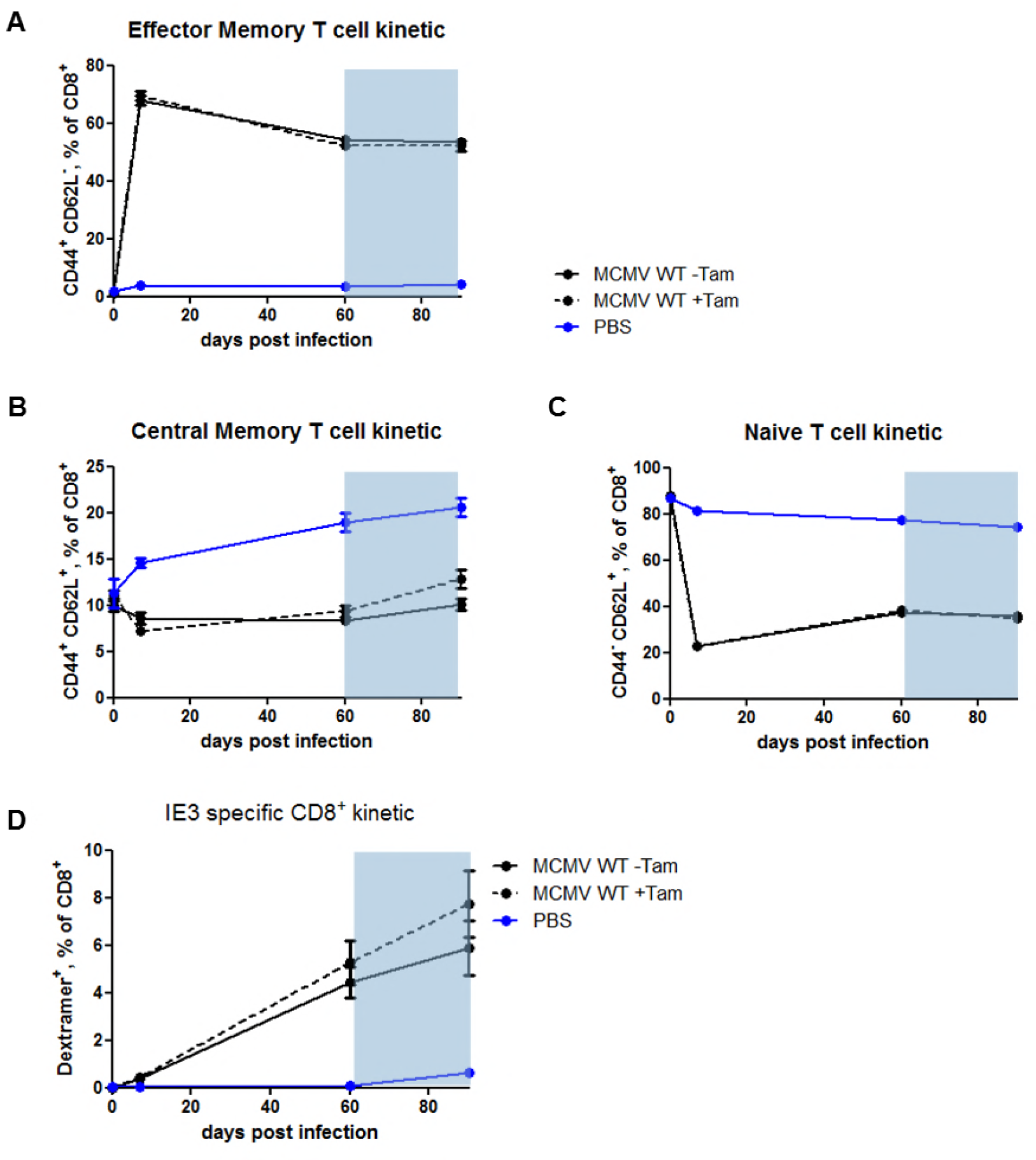
Tamoxifen treatment has no effect on any CD8 T-cell subsets in C67BL/6 wild-type animals. C57BL/6 wildtype mice were infected with 10^6^ PFU MCMV or 200 μl PBS (mock). At 60 dpi mice received food pellets containing 400 mg/kg Tamoxifen for 4 weeks (indicated with a grey rectangle). Blood was collected and analysed at 0, 7, 60, and 88 dpi. Blood leukocytes were stimulated with the IE3 peptide (RALEYKNL) and stained for CD8, CD44, CD62L, and IFNγ. (A) Kinetic of TEM (CD44+ CD62L^-^) CD8 T-cells before and after Tamoxifen treatment. (B) Kinetic of TCM (CD44^+^ CD62L^+^) CD8 T-cells befor and after Tamoxifen treatment. (C) Kinetic of naïve (CD44^-^ CD62L^+^) CD8 T-cells before and after Tamoxifen treatment. (D) Kinetic of IE3-specific CD8 T-cells before and after Tamoxifen treatment.

**Supplementary Figure 5:**
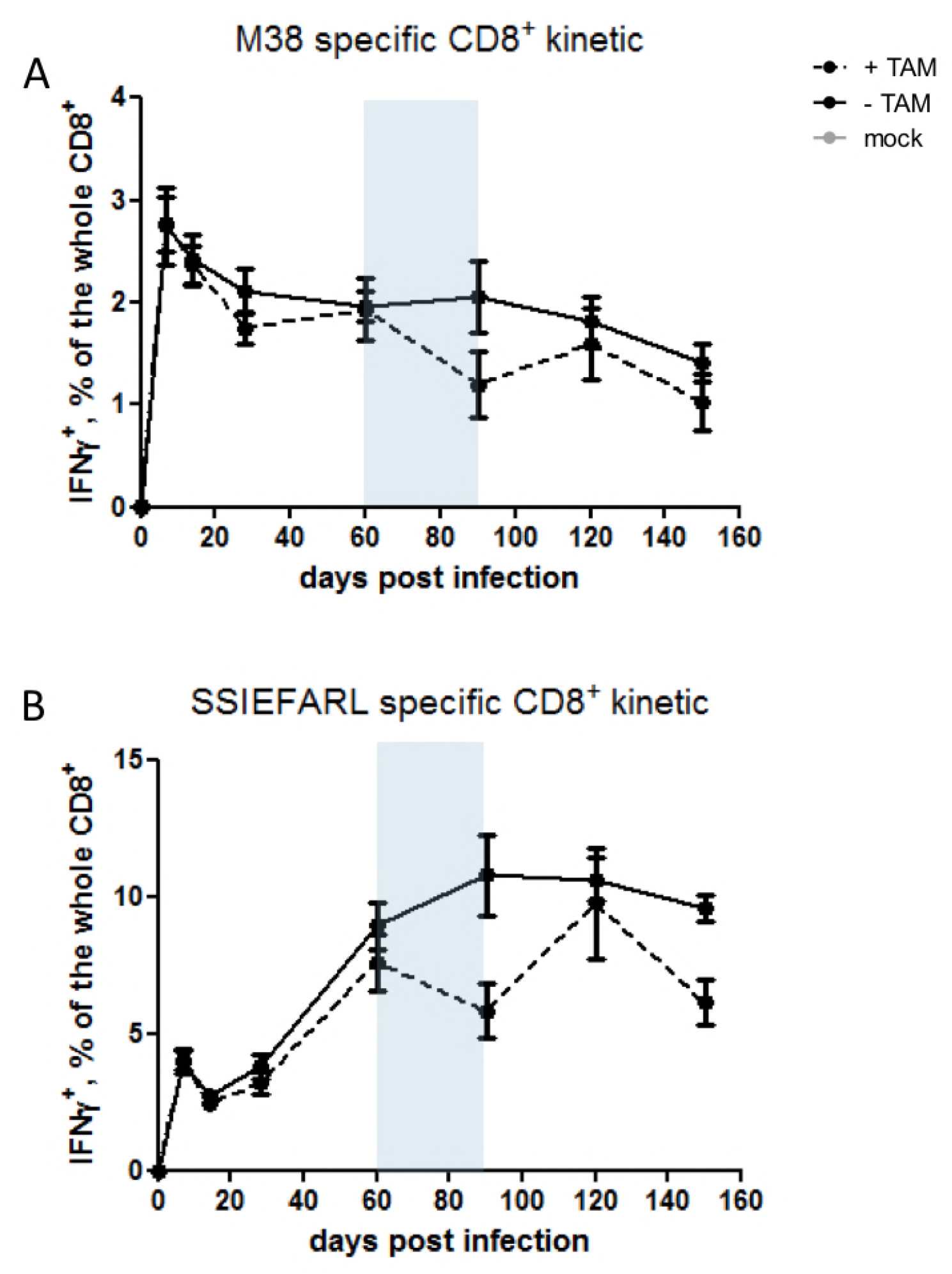
MCMV-specific CD8 T-cells are depleted by Tamoxifen treatment in R26 CreER^T2^ mice. (A, B) R26CreER^T2^ mice were infected with 10^6^ PFU MCMV or 200μl PBS (mock). 60 days post infection mice received food pellets with 400 mg/kg Tamoxifen for 4 weeks (indicated by the grey rectangle). Blood was collected and analysed at 0, 7, 14, 28, 60, 90, 120, 150, 180, and 270 dpi. Blood leukocytes were stimulated with the M38 peptide (SSPPMFRV) or the gB peptide (SSIEFARL) and stained for CD8, CD44, CD127, and IFNγ expression. (A) Kinetic of M38-specific CD8 T-cells before, during, and after Tamoxifen treatment. (B) Kinetic of gB-specific CD8 T-cells before, during, and after Tamoxifen treatment. Data are pooled from three independent experiments with up to 15 mice per group in total. Error bars indicate SEM. Significance was assessed by Mann-Whitney test. \*\**p* < 0.01. \*\*\*\**p* < 0.0001.

